# MCART1 is required for mitochondrial NAD transport

**DOI:** 10.1101/2020.08.28.267252

**Authors:** Nora Kory, Jelmi uit de Bos, Sanne van der Rijt, Nevena Jankovic, Miriam Güra, Nicholas Arp, Izabella A. Pena, Gyan Prakash, Sze Ham Chan, Tenzin Kunchok, Caroline A. Lewis, David M. Sabatini

## Abstract

The nicotinamide adenine dinucleotide (NAD+/NADH) pair is a cofactor in redox reactions and is particularly critical in mitochondria as it connects substrate oxidation by the tricarboxylic acid (TCA) cycle to ATP generation by the electron transport chain (ETC) and oxidative phosphorylation. While a mitochondrial NAD^+^ transporter has been identified in yeast, how NAD enters mitochondria in higher eukaryotes is unknown. Here, we mine gene essentiality data from human cell lines to identify *MCART1* (*SLC25A51*) as co-essential with ETC components. *MCART1*-null cells have large decreases in TCA cycle flux, mitochondrial respiration, ETC complex I activity, and mitochondrial levels of NAD^+^ and NADH. Isolated mitochondria from cells lacking or overexpressing *MCART1* have greatly decreased or increased NAD uptake in vitro, respectively. Moreover, *MCART1* and *NDT1*, a yeast mitochondrial NAD^+^ transporter, can functionally complement for each other. Thus, we propose that MCART1 is the long sought mitochondrial transporter for NAD in human cells.

## Introduction

Nicotinamide adenine dinucleotide (NAD) is an essential cofactor in redox metabolism and metabolic signaling. As cofactors of metabolic enzymes, NAD^+^ and its reduced form, NADH, function in redox reactions in central metabolic pathways including glycolysis, TCA cycle, oxidative phosphorylation, one-carbon metabolism and control the direction of flux through these pathways. As a cosubstrate of the sirtuins and poly-(ADP-ribose) polymerases, NAD^+^ mediates posttranslational modification of metabolic enzymes, DNA repair and chromatin-modifying proteins, and other factors involved in stress response and signaling pathways that connect metabolism to different physiological responses (*1*). NAD levels decline in aging and administration of NAD precursors is currently being tested in the clinic as a measure to prevent age-associated diseases ((*2, 3*); clinicaltrials.gov).

Despite its importance for many cellular processes, the compartmentalization of NAD synthesis and its transport remain poorly understood. Up to ~70% of cellular NAD has been reported to be present within mitochondria (*4*), where it serves as an electron carrier connecting fuel oxidation in the TCA cycle with the electron transport chain (ETC) and thus ATP synthesis. Strikingly, if and how NAD is imported into mitochondria in higher eukaryotes is not known. While mitochondrial and chloroplast NAD^+^ transporters have been identified in yeast and plants (*5, 6*), the functional ortholog in animals remains elusive.

### MCART1 is an inner mitochondrial membrane protein required for mitochondrial respiration

To identify novel genes required for mitochondrial respiration, we used a previously described approach (*7*) to mine the Achilles dataset containing gene essentiality scores from 341 cell lines (*8*) for genes that were co-essential with the nuclear-encoded core component of respiratory complex I of the ETC, *NDUFS1*. This analysis identified a cluster of ~400 co-essential genes, most of which were previously annotated as encoding mitochondrially-localized proteins, including components of or assembly factors for the ETC, and components of the mitochondrial translation machinery, whose main function is to synthesize the 13 mitochondrially-encoded components of the ETC (Fig. 1A). Embedded in this gene cluster, we identified a thus far unstudied gene, *MCART1* (*SLC25A51*) (Fig. 1A). The top 20 genes co-essential with *MCART1* are involved in ETC function or mitochondrial translation (Fig. 1B and fig. S1A), and this gene cluster is enriched specifically for genes that impact ETC function or related processes, such as the TCA cycle, but not other mitochondrial processes, such as mitophagy or mitochondrial fusion (Fig. 1C).

**Figure 1.**
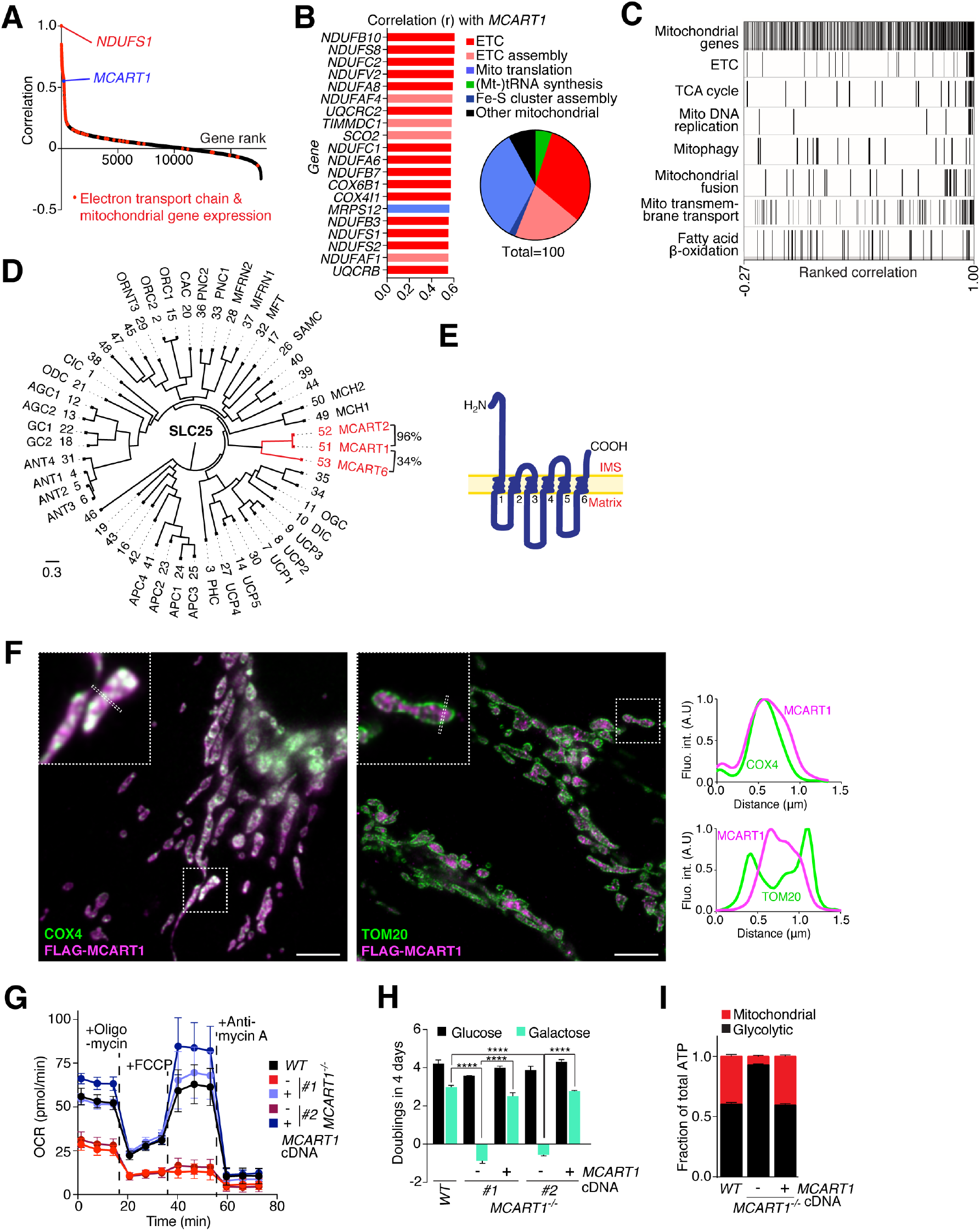
*MCART1* (*SLC25A51*) is an inner mitochondrial membrane solute carrier required for ETC function. **(A)** The respiratory complex I core subunit *NDUFS1* is co-essential with a cluster of genes involved in electron transport chain function. Gene essentiality scores from the Achilles dataset across 341 different cell lines (*8*) were correlated to identify genes co-essential with *NDUFS1* as described (*7*). Genes with a gene ontology annotation of electron transport chain or mitochondrial gene expression are highlighted in red. **(B)** *MCART1* is co-essential with genes involved in mitochondrial respiration. Gene ontology annotation of the top 20 (left panel) and top 100 (right panel) highest correlated genes with *MCART1*. ETC – electron transport chain; Mt – mitochondrial; Fe-S – ironsulfur. **(C)** Barcode plot displays enrichment of components of the electron transport chain, tricarboxylic acid (TCA) cycle and mitochondrial DNA replication, but not of other mitochondrial processes. All genes from the Achilles dataset are plotted from lowest to highest correlating with MCART1 and genes of the depicted category are displayed by black lines. **(D)** Phylogenetic tree of the human SLC25 family of mitochondrial carriers, which includes MCART1 and the closely related MCART2 and MCART6 in red. The number of each family member and, if they have one, their alias is included. % Sequence identity of MCART1, 2 and 6 is shown. **(E)** Model of the predicted topology of MCART1 in the mitochondrial inner membrane. Transmembrane helices are indicated by numbers. IMS – intermembrane space. **(F)** Super-resolution microscopy confirms MCART1 localization to the inner membrane of mitochondria. Wild-type HeLa cells transiently expressing FLAG-MCART1 were processed for immunofluorescence detection of the FLAG epitope (magenta) and the mitochondrial inner membrane marker cytochrome c oxidase subunit 4 (COX4) (left panel, green) or the outer mitochondrial membrane marker Tom20 (right panel, green) and imaged by STED microscopy. Overlap of magenta and green channels is shown in white. Scale bar is 2 μm. Line profiles show fluorescent signals of each channel across mitochondria where marked by the dotted rectangles in images. **(G)** Loss of MCART1 decreases the oxygen consumption rate (OCR) of cells. Oxygen consumption rate was measured by Seahorse Extracellular Flux Analysis (mean ± SD; *n* ≥ 13 technical replicates). First three time points are basal respiration, subsequent time points are with serial injections of oligomycin, FCCP, and antimycin A/rotenone, respectively. **(H)** *MCART1*-null cells are unable to proliferate using galactose as the main carbon source. Proliferation of wildtype and *MCART1*-null Jurkat cells was assayed in RPMI containing glucose or galactose as indicated (mean ± SD; *n* = 3; \*\*\**P* < 0.001, \*\*\*\**P* < 0.0001). **(I)** Mitochondrial ATP production is strongly reduced relative to glycolytic ATP production upon MCART1 loss (mean ± SD; *n* ≥ 13 technical replicates). Graphs were generated from data in the Seahorse experiment in fig. S2A using the Seahorse Report Generator.

MCART1 is an unstudied member of the SLC25 family of mitochondrial triple repeat carriers comprising 53 members in humans (Fig. 1D). Two other SLC25 family members, MCART2 (SLC25A52) and MCART6 (SLC25A53), are particularly similar to MCART1, with MCART2 having a striking 96% sequence identity (NCBI blast). However, while *MCART1* is expressed at considerable levels across tissues and commonly used cell lines, the expression of *MCART6* is generally lower and that of *MCART2* appears restricted to the testis (fig. S1, B and C). Like other members of the SLC25 family, the MCART1 protein is predicted to have six transmembrane domains, with both N- and C-termini facing the intermembrane space matrix based on recent proteomics proximity-labeling analysis (Fig. 1E; (*9*), topology analysis with Protter (*10*)). As expected, FLAG-tagged MCART1 co-localized with the inner mitochondrial membrane protein COX4 in HeLa cells (fig. S1C), and endogenous MCART1 was enriched in mitochondria purified from Jurkat cells (fig. S1D). Super-resolution microscopy revealed that MCART1 localizes to the inner mitochondrial membrane, consistent with a function in transporting a metabolite into mitochondria (Fig. 1F).

To test whether MCART1 has a role in oxidative phosphorylation, we deleted *MCART1* in human Jurkat leukemic T-cells using CRISPR-Cas9. Two clones with complete deletion of *MCART1* were chosen for downstream phenotypic analysis (fig. S1, E and F). *MCART1*-null cells had a strong proliferation defect in full media at early passages (fig. S1G), and we observed an increased acidification of the culture media despite their reduced proliferation suggesting a defect in mitochondrial function. Indeed, cells lacking MCART1 had a dramatically reduced oxygen consumption rate, stemming from reduced basal and maximal respiration, spare respiratory capacity, proton leak, and ATP production as measured by Seahorse extracellular flux analysis (Fig. 1G, fig. S1H). *MCART1*-null cells were also unable to proliferate in media containing galactose instead of glucose as a carbon source, conditions under which cells must generate ATP from mitochondrial respiration (Fig. 1H,). This phenotype was also observed in HeLa, HEK-293T, and 143B cells lacking *MCART1* (fig. S1P). Indeed, *MCART1*-null cells were defective in mitochondrial ATP production and instead generated the vast majority of their ATP from cytosolic glycolysis (Fig. 1I and fig. S1L). Importantly, re-expression of a guide-resistant cDNA for MCART1 reversed all of these defects (Fig. 1, G-H, and figs. S1, G, H, L). Expression of MCART2, the closest MCART1 homolog, but not of MCART6, rescued proliferation on galactose (fig. S1I-K). Interestingly, the proliferation defect of *MCART1*-null cells was not rescued by the addition to the media of metabolites known to bypass different aspects of mitochondrial function, such as pyruvate and uridine, formate, hypoxanthine-thymidine, or aspartate even in cells expressing a plasma membrane aspartate transporter (*11-16*)(fig. S1, N and O). The inability of these metabolites to rescue the proliferation of *MCART1*-null cells suggests that ATP levels are limiting for proliferation, as described previously for mitochondrial dysfunction caused by loss of ETC components (*17*), and we did indeed observe a slight decrease in total cellular ATP levels (fig. S1M).

### Loss of MCART1 results in defects in mitochondrial metabolism and ETC complex I activity without affecting mitochondrial integrity

Mitochondrial dysfunction and respiratory defects are often due to defects in mitochondrial replication, translation or structural integrity that lead to loss of respiratory chain complexes. However, loss of MCART1 did not change mitochondrial or cristae morphology nor mitochondrial DNA or mass (fig. S2, A-D). Furthermore, the mitochondrial membrane potential and levels of mitochondrially as well nuclear-encoded mitochondrial proteins were only marginally, if at all, affected (fig. S2, E and F). To test whether the ETC generally or a specific respiratory complex was affected in *MCART1*-null cells, we measured the activity of the respiratory chain by providing (artificial) substrates for each respiratory chain complex to permeabilized cells and analyzed oxygen consumption rate by Seahorse extracellular flux analysis. Remarkably, when complex I substrates were provided oxygen consumption was ablated (and respiratory control ratio decreased) in *MCART1*-null cells, while the oxygen consumption rate was comparable to that in wild-type cells, or cells re-expressing *MCART1* when substrates for other respiratory chain complexes were provided, such as succinate feeding into complex II (Fig. 2, A and B, fig. S2, G, H, I). When we added substrates directly to mitochondrial lysates, the NADH:ubiquinone oxidoreductase activity of complex I in *MCART1*-null cells did not differ from that in wild-type cells, arguing that complex I levels, assembly, or function itself were not perturbed (Fig. 2C). Collectively, these data suggested that the mitochondrial dysfunction caused by MCART1 deletion was most likely caused by loss of a metabolite in the mitochondria with a specific role in complex I of the ETC.

**Figure 2.**
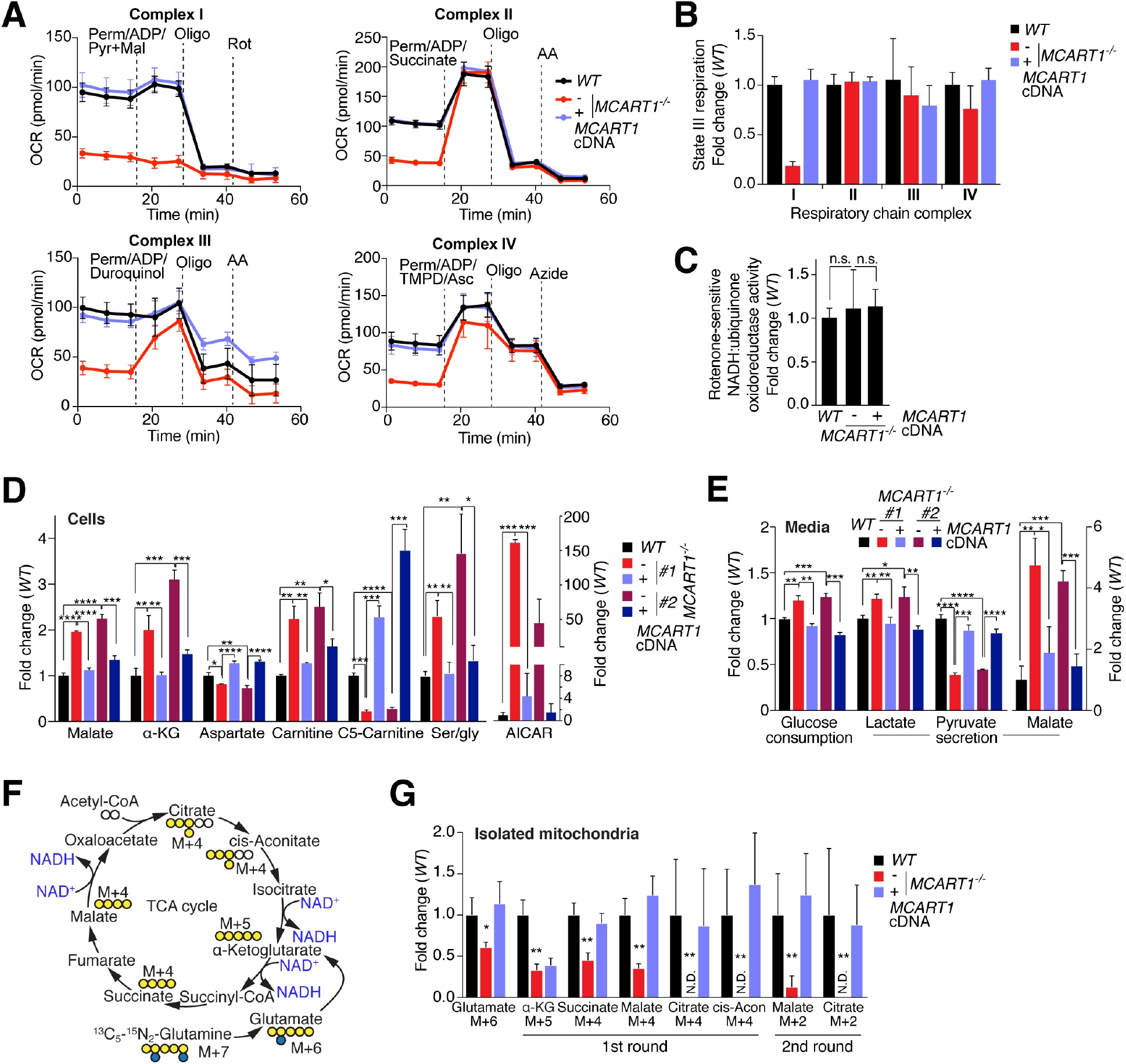
Loss of MCART1 causes loss of ETC complex I activity and defects in mitochondrial metabolism without affecting mitochondrial integrity. **(A)** Loss of MCART1 diminishes the activity of respiratory complex I, but not other complexes. Oxygen consumption rate (OCR) of indicated cells permeabilized and supplemented with ADP and complex I-IV substrates was measured by Seahorse extracellular flux analysis (mean ± SD; *n* = 3 technical replicates). Mal – malate; perm – permeabilizer; pyr – pyruvate; rot – rotenone; AA – antimycin A. **(B)** Loss of MCART1 diminishes complex I-dependent state 3 respiration. Graph was calculated from the data in Fig. 2B (mean ± SD; *n* = 3 technical replicates; data from the second *MCART1-null* clone is shown in fig. S2I). **(C)** Rotenone-sensitive NADH:ubiquinone activity in mitochondrial lysates is not dependent on MCART1 indicating complex I is functional in *MCART1*-null cells. (mean ± SD; *n* = 3; n.s. – not significant). **(D)** Several mitochondrial and mitochondria-derived metabolites are changed in cells lacking MCART1 compared to their wild-type counterparts. Metabolites were measured by LC-MS in extracts from indicated cells (mean ± SD; *n* = 3; *\*\*P* < 0.01). α-KG – α-ketoglutarate; ser – serine; gly – glycine; AICAR – 5-Aminoimidazole-4-carboxamide ribonucleotide. **(E)** Loss of MCART1 increases glucose consumption and lactate and malate excretion, and decreases pyruvate secretion. Medium metabolites were extracted after growing cells for 48 hours in RPMI media (mean ± SD; *n* = 3). Values are normalized to cell number. **(F)** Glutamine tracing scheme used to measure TCA cycle flux. **(G)** TCA cycle flux depends on MCART1. Isolated mitochondria were incubated with ^13^C_5_,^15^N_2_-glutamine, malate and ADP, and the metabolites generated from labeled glutamine in the first (α-KG M+5, succinate/malate/citrate/cis-aconitate M+4) and second rounds (malate/citrate M+2) of the TCA cycle according to the tracing scheme in the left panel were detected by LC-MS. (mean ± SD; *n* = 3) α-KG – α-Ketoglutarate; cis-Acon – cis-aconitate.

To understand how mitochondrial metabolism was altered upon loss of MCART1, we turned to LC-MS-based metabolomics analyses. A number of mitochondria-derived metabolites, such as TCA cycle intermediates, the fatty acid carrier carnitine, the branched chain amino acid breakdown product C5-carnitine, and the purine synthesis intermediate 5-Aminoimidazole-4-carboxamide ribonucleotide (AICAR), were dramatically changed in *MCART1*-null cells (Fig. 2D). *MCART1*-null cells also consumed glucose and secreted lactate and malate at higher rates than wild-type cells, while the observed net secretion of pyruvate from cells to the media was decreased (Fig. 2E). These changes in intracellular and media metabolites indicated extensive alterations in TCA cycle and one-carbon flux, fatty acid/branched chain amino acid-oxidation and ETC function, all of which are major mitochondrial metabolic pathways, and are consistent with a switch to aerobic glycolysis in *MCART1*-null cells.

To assess the capacity for mitochondria to catabolize nutrients, we probed TCA cycle flux in mitochondria specifically by incubating isolated mitochondria with stably isotope-labeled ^13^C_5_-^15^N_2_-glutamine (M+7) and measuring newly synthesized (labeled) TCA cycle metabolites by LC-MS (Fig. 2F). The use of isolated mitochondria in this assay is critical because some TCA cycle intermediates can also be synthesized in the cytosol or nucleus (*18-20*). Despite significant glutaminolysis still occurring as assessed by M+6 glutamate levels, detectable TCA cycle intermediates produced in the first (M+5, M+4) and second rounds (M+2) of the TCA cycle were dramatically decreased or undetectable when *MCART1*-null mitochondria were used in the reaction (Fig. 2G). Of note, when glutamine tracing was performed in whole cells no decrease in the production of TCA cycle metabolites was observed indicating an increased activity, perhaps due to compensation, of cytosolic isoenzymes (fig. S2J).

### Depletion of NAD^+^ and NADH in mitochondria from *MCART1*-null cells

To understand how loss of MCART1 affects metabolism in mitochondria specifically, we isolated mitochondria using the Mito-IP approach and broadly profiled metabolites by LC-MS, allowing us to detect the mitochondrial metabolites whose levels most dramatically change upon loss of MCART1 (*21*). Strikingly, the largest difference between mitochondria with and without MCART1 was in the dinucleotide NAD in both its oxidized (NAD^+^) as well as reduced form (NADH; Fig. 3A). NAD^+^ and NADH were greatly diminished in mitochondria of *MCART1*-null cells, while their whole cell levels were unaffected (Fig. 3B, fig. S3A,B). We also observed decreases in the TCA cycle intermediates cis-aconitate, alpha-ketoglutarate, and malate specifically in mitochondria, and an overall decrease in phosphoenolpyruvate, which is generated from the TCA cycle intermediate oxaloacetate, consistent with our TCA cycle flux analysis. Glutamate was increased, consistent with repressed TCA cycle anaplerosis, and ATP, ADP, and other mitochondrial metabolites were only slightly or not significantly changed (Fig. 3B, fig. S3A,B).

**Figure 3.**
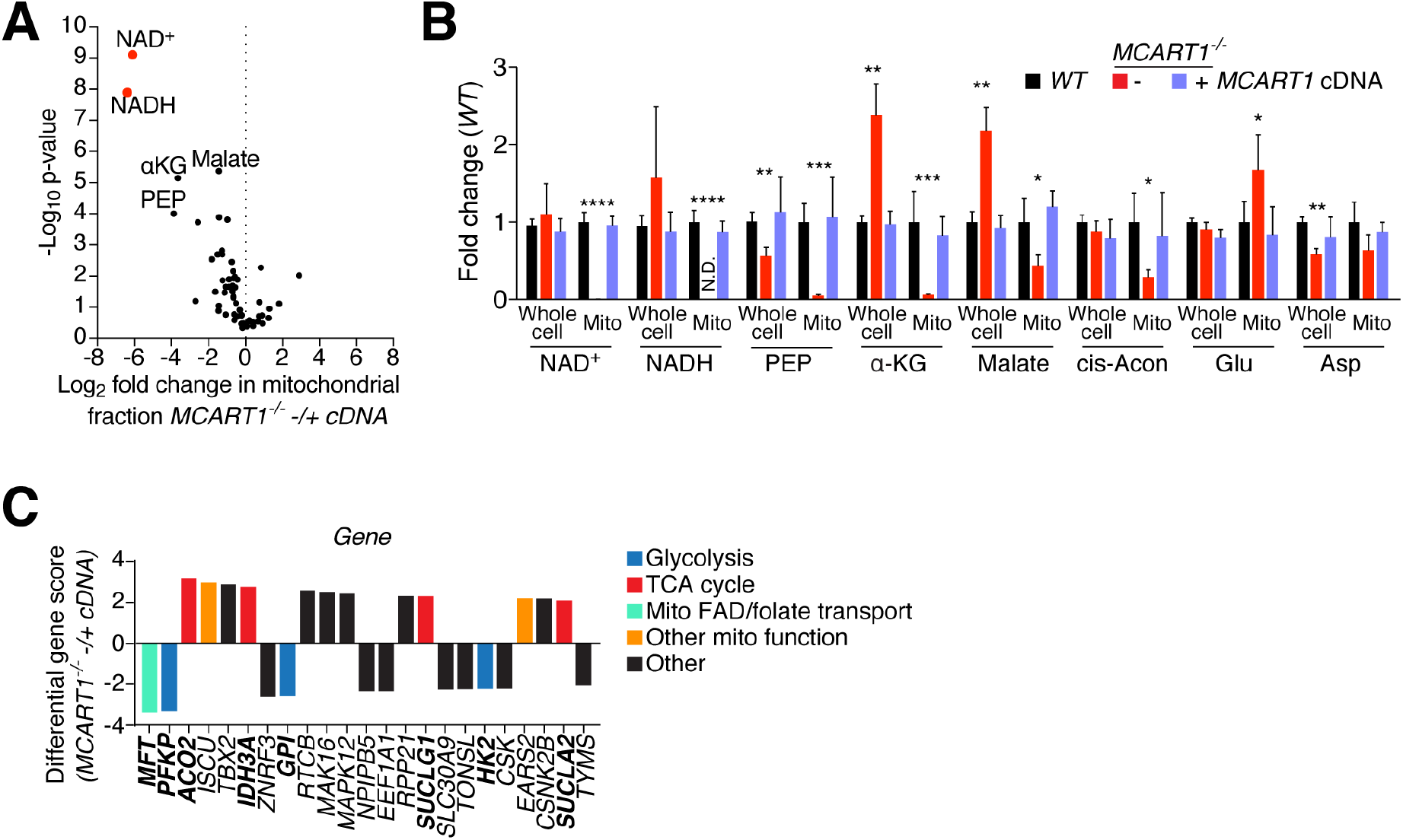
NAD^+^ and NAD are depleted in the mitochondria of *MCART1-null* cells. **(A)** NAD^+^ and NADH are the most depleted metabolites in mitochondria of *MCART1-null* cells. The log2 fold change of metabolites detected in mitochondria isolated from *MCART1-null* cells versus in mitochondria from null cells expressing MCART1 cDNA (mean; *n* = 3). **(B)** Loss of MCART1 depletes NAD^+^ and NADH in mitochondria and reduces TCA cycle intermediates. Whole cell and mitochondrial (mito) metabolite levels in indicated cells were measured by LC-MS using the Mito-IP method, data from two independent experiments were combined (mean ± SD; *n* > 5). Asterisks denote statistically significant differences of *MCART1-null* samples with both wild-type cells and cells re-expressing the *MCART1* cDNA (\**P* < 0.05, \*\**P* < 0.01, \*\*\**P* < 0.001, \*\*\*\**P* < 0.0001). PEP – phosphoenolpyruvate; α-KG – α-Ketoglutarate; cis-Acon – cis-aconitate; Glu – glutamate; Asp – aspartate. Data from the second *MCART1-null* clone is shown in fig. S2I. **(C)** *MCART1*-null cells depend on glycolytic enzymes and mitochondrial FAD/folate transporter. Top-scoring genes from the MCART1 synthetic lethality screen. Genes were ranked according to the differential gene score in *MCART1*-null versus control cells. (Mito – mitochondrial; TCA – tricarboxylic acid; FAD – flavin adenine dinucleotide).

The NAD^+^/NADH redox pair is a critical cofactor in mitochondrial metabolism and acts as an electron carrier feeding into ETC complex I. Thus, the specific loss of complex I activity and the metabolite changes observed in *MCART1*-null cells are consistent with a loss of mitochondrial NAD and together these results suggested MCART1 functions in the uptake of NAD or an NAD precursor into mitochondria. In subsequent metabolite profiling experiments we therefore used a metabolite extraction method optimized for preserving NAD-related metabolites (adapted from (*22*)).

To corroborate this possible function of MCART1 in an unbiased way we identified genes synthetically lethal with *MCART1* using a negative selection-CRISPR-Cas9 screen. Consistent with our previous results, several genes involved in glycolysis (which uses NAD in the cytosol) were selectively essential in cells lacking MCART1, while control cells re-expressing the MCART1 cDNA depended more on TCA cycle enzymes (which use NAD in mitochondria) (Fig. 3C, S3C). Notably, the mitochondrial folate carrier MFT/SCL25A32, which has been proposed to transport the redox cofactor FAD and folates into mitochondria (*23-27*), was the most selectively essential gene in *MCART1*-null cells. This could be explained by the notion that depletion of one redox cofactor (NAD) from mitochondria upon *MCART1* loss results in increased dependence on another (FAD).

### A yeast mitochondrial NAD^+^ transporter but not a predicted substrate-binding mutant of MCART1 rescues loss of MCART1

In mammals, de novo synthesis of NAD from tryptophan occurs primarily in the liver, while most other tissues rely on NAD synthesis or salvage from its precursors niacin, nicotinamide (Nam), nicotinamide riboside (NR) or nicotinamide mononucleotide (NMN) (*28, 29*). The precise step at which NAD is transported into mitochondria and whether NAD can be synthesized in mitochondria is unclear (Fig. 4A). Both NMN and NAD have been proposed to be transported into mitochondria of human cells (*28, 30, 31*). However, deletion of NMNAT3, the mitochondrial NAD synthesis isoenzyme that uses NMN to generate NAD, which is expressed in Jurkat cells, did not affect respiration or mitochondrial NAD levels making NMN as the transported species highly unlikely (fig. S4A-F).

**Figure 4.**
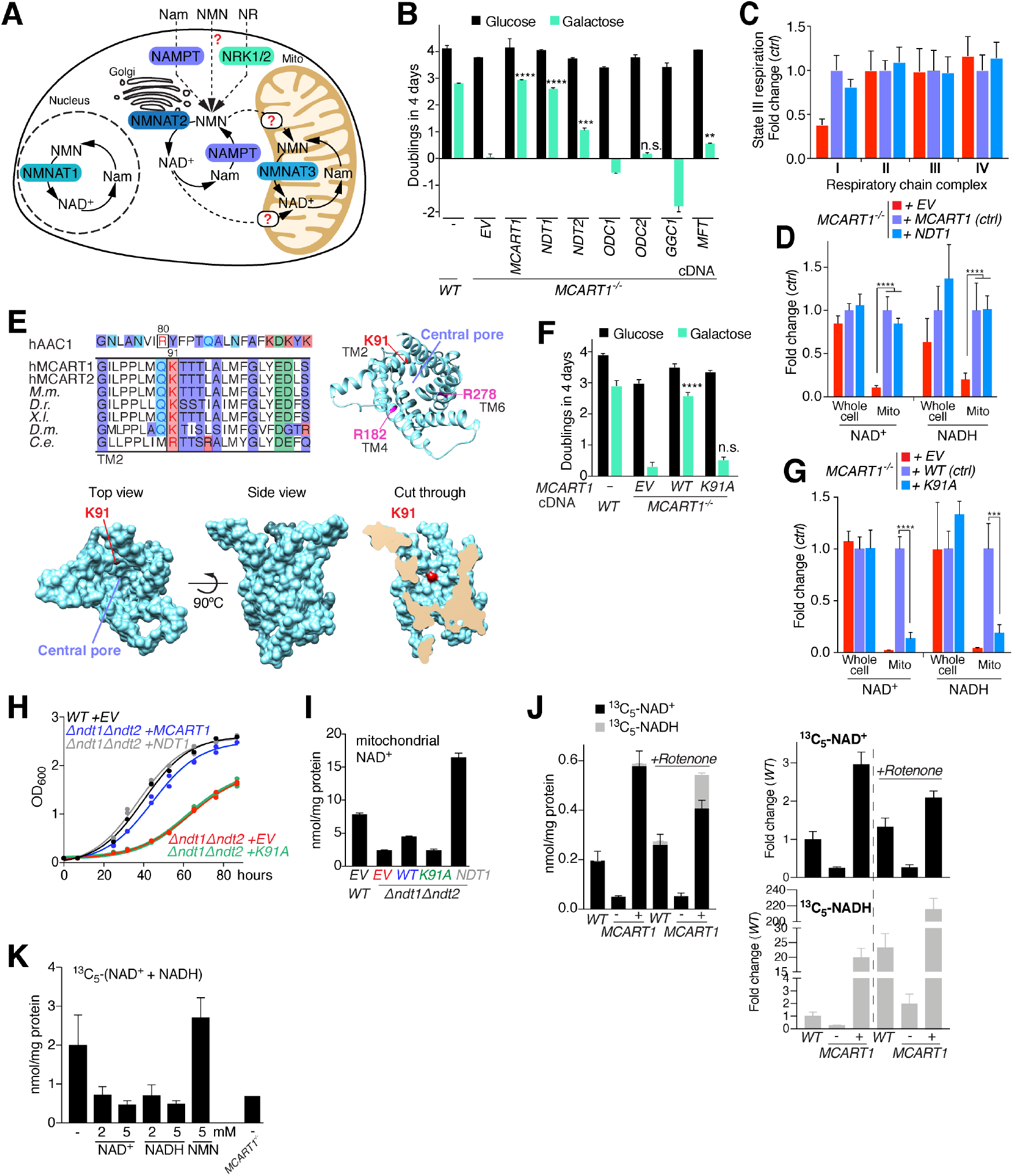
A yeast mitochondrial NAD^+^ transporter rescues the defects of *MCART1-null* cells and mitochondrial NAD transport depends on MCART1. **(A)** Schematic of the NAD salvage pathway. (Nam – nicotinamide; NMN – nicotinamide mononucleotide; NRK – nicotinamide riboside kinase; NAMPT – nicotinamide phosphoribosyltransferase; NMNAT – nicotinamide-nucleotide adenylyltransferase). **(B)** The yeast mitochondrial NAD^+^ transporter NDT1 but not close sequence homologs of MCART1 rescue mitochondrial respiration as determined by growth in galactose as the carbon source. Single-cell-derived knockout Jurkat cells were transduced with an empty vector (EV) or cDNAs of MCART1, the mitochondrial FAD/folate carrier (MFT) or yeast mitochondrial transporters. Asterisks denote statistically significant differences in proliferation in media containing galactose as the carbon source between the cells expressing the empty vector and the solute carrier homologs (mean ± SD; *n* = 3; \*\**P* < 0.01, \*\*\**P* < 0.001, \*\*\*\**P* < 0.0001). N.s. – not significant; ODC – oxodicarboxylate carrier, GGC – GTP/GDP carrier. **(C)** The yeast mitochondrial NAD^+^ transporter NDT1 rescues complex I activity in *MCART1*-null cells (mean ± SD; *n* > 3 technical replicates). EV – empty vector. **(D)** The yeast mitochondrial NAD^+^ transporter NDT1 rescues mitochondrial NAD levels in *MCART1*-null cells (mean ± SD; *n* > 3; \*\*\*\**P* < 0.0001). EV – empty vector. **(E)** Sequence alignment of MCART1 homologs and sequence comparison with the ADP/ATP carrier identify lysine residue 91 in transmembrane domain 2 as a potential substrate contact point. Arginines 182 and 278 are potential other substrate contact points. TM – transmembrane domain. **(F)** Mutation of lysine 91 to alanine abolishes the ability of MCART1 to rescue growth on galactose. *MCART1*-null cells infected with wild-type MCART1 cDNA serve as control cells. (mean ± SD; *n* = 3; \*\*\*\**P* < 0.0001). EV – empty vector. **(G)** MCART1 K91A does not rescue mitochondrial NAD^+^ and NADH levels. *MCART1*-null cells infected with wild-type MCART1 cDNA serve as control cells (mean ± SD; *n* = 4; \*\*\**P* < 0.001, \*\*\*\**P* < 0.0001). **(H)** MCART1 but not the point mutant MCART1K91A rescues proliferation of *ndt1Δndt2Δ S. cerevisiae*. Yeast were inoculated at an OD_600_ of 0.1 in synthetic minimal media with 2% ethanol. Two independent clones are shown at each time point with the exception of wild-type cells where one clone is shown. EV – empty vector. **(I)** Mitochondrial NAD^+^ concentrations in wild-type *S. cerevisiae*, or *ndt1Δndt2Δ* yeast transformed with empty vector (EV), MCART1 or MCART1K91A normalized to mitochondrial protein content. NADH was not detected (mean ± SD; *n* = 3 technical replicates). **(J)** Transport of NAD^+^ into mitochondria depends on MCART1. Mitochondria purified from wild-type, *MCART1-null*, or *MCART1-null* cells overexpressing the MCART1 cDNA were incubated with 50 μM stably isotope labeled ^13^C_5_-NAD^+^ for 10 min at 30 °C with or without 5 μM rotenone. Levels of uptaken ^13^C_5_-NAD^+^ and generated ^13^C_5_-NADH were quantified by LC-MS. Separate plots for the fold change of NAD^+^ and NADH are shown on the right. (mean ± SD; *n* = 3 uptake reaction replicates). The data are representative of three independent experiments. **(K)** MCART1-dependent mitochondrial NAD transport is competed by NAD^+^ and NADH but not the NAD synthesis intermediate NMN. Wild-type mitochondria were incubated for 10 min with 500 μM ^13^C_5_-NAD^+^ in the presence of unlabeled metabolites as indicated. The level of background binding/uptake in *MCART1-null* mitochondria is shown on the right. The sum of quantified ^13^C_5_-NAD^+^ and ^13^C_5_-NADH is shown (mean ± SD; *n* = 3 uptake reaction replicates). The data are representative of two independent experiments.

Based on sequence alignment and previously published mutational and structural analysis of the ADP/ATP carrier (*32-35*) we were able to model the structure of MCART1. We identified three residues conserved across species, lysine 91, arginine 182, and arginine 278, located on the inside of the pore as potential substrate contact points (Fig. 4E). Mutation of the conserved lysine residue 91, which is part of transmembrane helix 2, or arginine 278, part of transmembrane helix 6, to alanine abolished the ability of MCART1 to rescue mitochondrial NAD levels and respiration (Fig. 4, F and G, fig. S4G) without significantly affecting protein stability or localization (fig. S4, I and J), indicating that MCART1 is likely a transport channel and not simply an auxiliary factor for a transporter.

While a mammalian mitochondrial NAD transporter has not been identified, NDT1 and NDT2 are well known as the mitochondrial NAD^+^ transporters in yeast and plants (*5, 6, 36, 37*). Like MCART1, NDT1 and 2 are part of the SLC25 family of mitochondrial transporters, but they are not closely related to MCART1 in sequence or predicted structure and sequence analysis has failed to reveal the mitochondrial NAD transporter in higher eukaryotes (*38*). To test whether NDT1 and NDT2 could functionally complement loss of MCART1 as would be predicted if MCART1 is required for NAD transport into mitochondria, we expressed in *MCART1*-null cells cDNAs for NDT1 or 2 codon-optimized for expression in human cells. NDT1 and 2 but neither of the closest yeast MCART1 homologs predicted based on sequence or structure, the 2-oxodicarboxylate carriers ODC1 and 2 and the GTP/GDP carrier GGC1, rescued mitochondrial respiration in *MCART1*-null cells (Fig. 4B, fig. S4, H and J). Importantly, NDT1 also rescued complex I activity in *MCART1*-null cells and mitochondrial NAD levels (Fig. 4,C and D). The lesser rescue ability of NDT2 is likely due to its lower expression/protein stability and activity (fig. S4H, (*5*)). Of note, overexpression of the closest homolog of NDT1 in humans, the mitochondrial FAD and folate transporter MFT/SLC25A32, which scored as selectively essential in *MCART1*-null cells in our CRISPR screen, only very slightly improved respiration in *MCART1*-null cells (Fig. 4B).

Having identified a putative functional ortholog of the yeast mitochondrial NAD transporters in mammalian cells we tested whether expression of MCART1 was able to rescue the phenotypes of yeast lacking the known mitochondrial NAD transporters NDT1 and NDT2. Indeed, MCART1, but not its K91A mutant, rescued the proliferation and mitochondrial NAD^+^ levels of *ndt1Δndt2Δ* yeast grown on minimal ethanol media (Fig. 4, H and I). MCART1 rescued proliferation nearly completely but NAD^+^ levels only partially, consistent with previous results suggesting a threshold exists below which mitochondrial NAD levels become limiting for proliferation (NADH was not detected, which is consistent with its low abundance in yeast mitochondria)(*5*).

### MCART1 is required for mitochondria to take up NAD

To test whether MCART1 was required for mitochondrial NAD uptake in mammalian cells, we isolated mitochondria from wild-type cells, *MCART1*-null cells, or the *MCART1*-null cells overexpressing the *MCART1* cDNA, incubated them in vitro with stable isotope-labeled ^13^C_5_-NAD^+^,and after 10 min quantified metabolites taken up and generated by the mitochondria by LC-MS. NAD uptake into mitochondrial lacking MCART1 was highly reduced and uptake into mitochondria from cells over-expressing MCART1 was increased compared to wild-type mitochondria (Fig.4J). In addition, loss of MCART1 greatly reduced, and its overexpression increased, the generation of labelled NADH in the reaction. Consistent with the oxidation of NAD^+^ into NADH by complex I requiring its transport into the mitochondrial matrix, the addition to the reaction of rotenone, an inhibitor of complex I-mediated NAD^+^ oxidation, increased the ratio of labeled NADH to NAD^+^. These results indicate that NAD^+^ was taken up in an MCART1-dependent manner, into the mitochondrial matrix, where it was enzymatically reduced (Fig. 4J, fig. S4K). NAD uptake remained dependent on MCART1 even upon the addition of succinate and ADP (fig. S4K), conditions under which the respiration of *MCART1*-null mitochondria is comparable to that of wild-type mitochondria (Figure 2A,B). Succinate/ADP had a membrane-potential-dependent effect on increasing the NADH to NAD^+^ ratio, presumably through reverse electron flux through complex I (fig. S4K). The MCART1-dependent uptake of NAD^+^ into mitochondria was competed by the addition of unlabeled NAD^+^ or NADH but not NMN (Fig. 4K).

## Discussion

Our results show that MCART1 is required for NAD uptake into mitochondria and thus the function of complex I in the electron transport chain. Loss of MCART1 and mitochondrial NAD leads to diminished flux through pathways relying on NAD in mitochondria such as the TCA cycle, mitochondrial one-carbon metabolism, and oxidative phosphorylation. Our work identifies for the first time a putative mitochondrial NAD importer in higher eukaryotes and thus addresses a long-standing question in the metabolism and mitochondrial biology fields. Further studies made possible by our identification of MCART1 including biochemical assays will reveal the precise biochemical properties of MCART1-dependent transport of NAD into mitochondria. Our findings corroborate a recent study claiming NAD itself is the imported species (*31*). In agreement, deletion of NMNAT3, the mitochondrial isoform of the NAD synthesis enzyme NMNAT previously implicated in maintenance of mitochondrial NAD levels (*28, 39*) did not affect mitochondrial NAD levels in our system or in several mouse tissues in a recent study (*40*). However, it is possible that there are other mitochondrial NAD synthesis enzymes and we do not exclude a contribution from the mitochondrial matrix to NAD synthesis and do not claim that transport from the cytosol is the only source of mitochondrial NAD in any eukaryotic cell type.

Our findings support that the mechanism of mitochondrial NAD transport and maintenance in mammals is related to that of yeast or plants, where mitochondrial NAD transporters have previously been identified (*5, 6*). Indeed, we find that MCART1 and the yeast NAD^+^ transporter NDT1 can functionally complement each other, although they are not homologous based on sequence. Several substrates have been reported for yeast and plant NAD transporters in addition to NAD^+^, such as AMP, GMP, FMN and FAD. Moreover, expression of the Arabidopsis thaliana mitochondrial NAD^+^ transporter NDT2 in HEK-293 cells resulted in dramatic growth retardation and a metabolic shift from oxidative phosphorylation to glycolysis, despite elevated mitochondrial NAD levels, indicating that the yeast and plant NDTs, particular NDT2, likely have other substrates besides NAD and different biochemical properties or regulation than mammalian MCART1 (*38*).

Further experiments, particularly with the purified protein reconstituted into proteoliposomes, will reveal whether MCART1 contains the NAD transport channel and can transport the additional substrates reported for Ndt1 and Ndt2. That overexpression of MCART1 is sufficient to increase the transport of NAD into isolated mitochondria suggests that MCART1 itself contains the NAD transport channel but we cannot exclude the possibility that other non-limiting proteins are necessary for MCART1-mediated NAD transport. In vitro transport assays combined with structural information will likely be necessary to understand the precise mechanism of NAD transport and how MCART1 activity and transport of NAD and related metabolites is regulated in order to not disrupt the redox balance between mitochondria and the cytosol. For example, MCART1 activity must not interfere with the malate-aspartate shuttle, which is responsible for exchanging reducing equivalents in the form of NADH across the inner mitochondrial membrane serving as an electron transport system for coupling NADH production by glycolysis in the cytosol to oxidative phosphorylation in mitochondria. Consistent with this, based on our uptake experiments, MCART1 likely has a high Km, which would support net transport only at high NAD concentrations or in situations when mitochondrial NAD actually becomes significantly depleted, such as in cells consuming mitochondrial NAD through sirtuins.

Despite displaying severe metabolic changes, and strong mitochondrial respiration and growth defects, *MCART1* is not absolutely cell essential as we were able to generate *MCART1*-null cells. Glycolysis provides ATP in *MCART1*-null cells and it is possible that NAD-requiring reactions shift to other cellular compartments where isoenzymes are present to compensate for loss of the production of certain metabolites in mitochondria. It is also possible that another transporter is able to maintain basal levels of NAD in mitochondria in the absence of MCART1. Candidates are the mitochondrial FAD/folate transporter MFT/SLC25A32, which was synthetically lethal with *MCART1* in our screen, as well as MCART2, although we could not detect its expression in our cell system. MCART1 has previously been reported to interact with assembly factors for iron-sulfur proteins and complex V (*41*), and thus it is possible that MCART1 affects mitochondrial respiration and metabolism in other ways than by maintaining mitochondrial NAD levels.

Determining how *MCART1*-null cells adapt and rewire their metabolism to loss of mitochondrial NAD could uncover metabolite or genetic interventions that are able to bypass MCART1 or electron transport chain function and might be useful strategies to treat complex I deficiency. In this way, MCART1 deficiency could be useful as a model for mitochondrial disease.

NAD plays a critical role in cellular and mitochondrial metabolism beyond its role as an enzymatic co-factor in redox reactions as a co-substrate of sirtuins and poly-ADP-ribose-polymerases. Three sirtuin homologs, SIRT3-5, serving as signaling factors connecting metabolism to cell state are present in mitochondria and their activity is likely coupled to mitochondrial NAD levels (*36, 42*). As NAD levels decline in aging and recent efforts are aimed at boosting cellular NAD levels to delay the onset of aging and age-related diseases (*1, 2, 43*), MCART1 emerges as a tool to study the role of the mitochondrial NAD pool in the regulation of these processes and perhaps as an interesting target to modulate life span.

## Acknowledgements

We thank all members of the Sabatini lab for helpful insights, suggestions and discussion, in particular Jessica Spinelli and Evgeni Frenkel, as well as Edmund Kunji at Cambridge University. We thank Heather Keys from the Functional Genomics Platform for technical assistance and helpful suggestions, as well as Nicki Watson and Wendy Salmon from the W.M. Keck Microscopy Facility at the Whitehead Institute and the Whitehead Institute FACS facility for technical assistance. We also thank Christine Hanko and Daniel Tom from Leica for help with STED super-resolution microscopy. Wild-type and *ndt1Δndt2Δ* yeast strains were a generous gift from Dr. Marina Vai. This work was supported by an HHMI Damon-Runyon Cancer Research Foundation fellowship and a NIH K99 award (CA241332) to N.K and by grants from the NIH to D.M.S (R01 CA103866, R01 CA129105, and R37 AI47389), the Department of Defense (W81XWH-07-0448), and the St. Baldrick’s Foundation. D.M.S. is an investigator of the Howard Hughes Medical Institute and an ACS Research Professor. N.K and D.M.S. are inventors on a patent application filed by Whitehead Institute relating to work described in this paper.

## Author Contributions

N.K. initiated the project, devised the research plan, designed and analyzed most experiments, and interpreted experimental results with guidance from D.M.S. N.K., J.u.d.B., S.v.d.R., N.J. and M.G. performed experiments with assistance from N.A. and G.P. N.K. wrote and D.M.S. edited the manuscript. I.A.P., C.A.L., S.H. C. and T.K. assisted with LC-MS sample preparation and ran the samples.

## Data and materials availability

All data are available in the manuscript or the supplementary material. All expression plasmids were deposited at Addgene.

## Supplementary Figures

**Figure S1.**
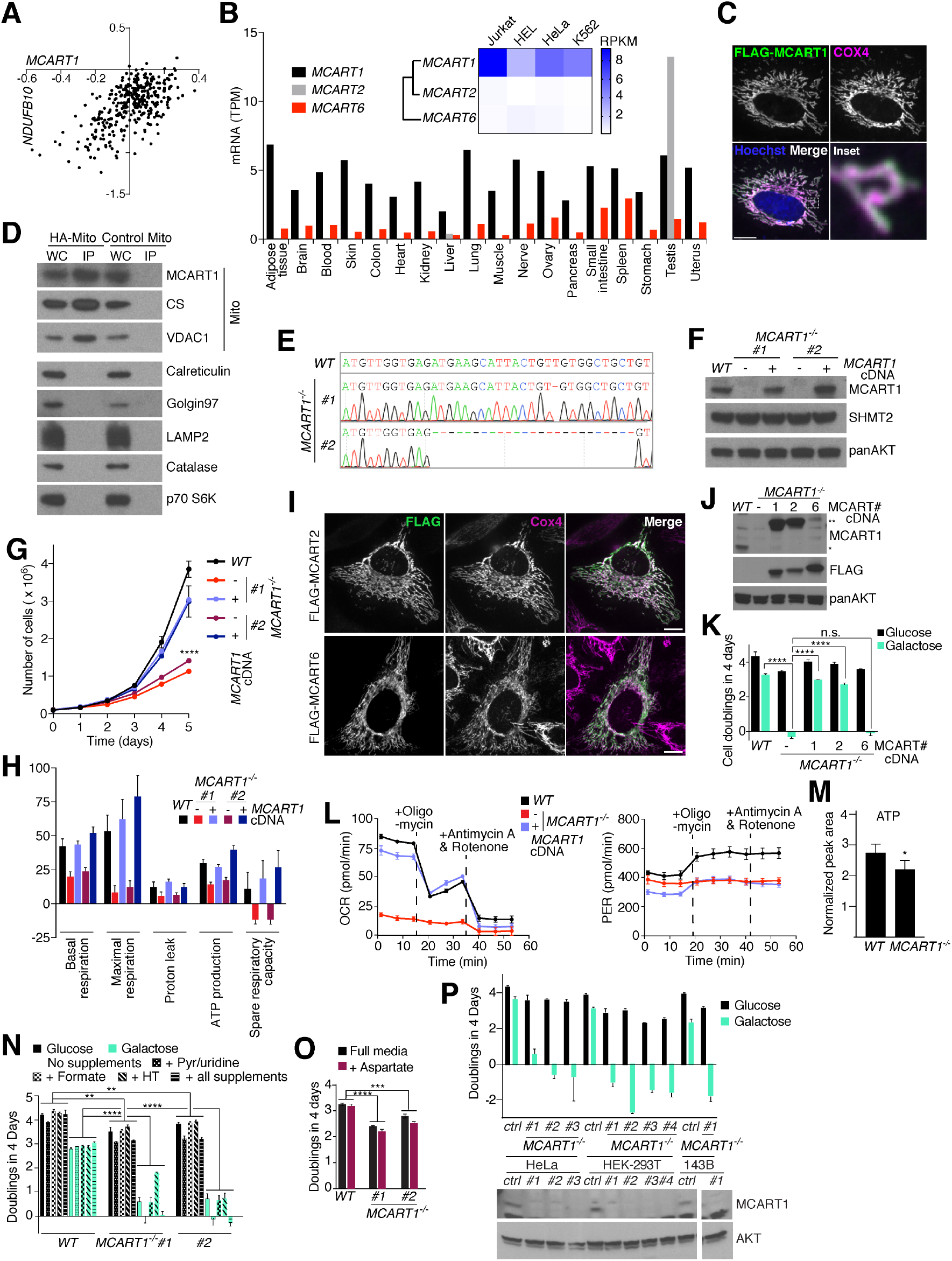
**(A)** *MCART1* correlates most strongly with *NDUFB10* in the Achilles dataset. Plotted are gene essentiality scores for *MCART1* and *NDUFB10* over a panel of 341 cancer cell lines (*8*). **(B)** mRNA levels of human *MCART* homologs in commonly used cell lines and normal tissues. RPKM (Reads Per Kilobase Million) levels were extracted from the Cancer Cell Line Encyclopedia (*56*) and TPM (Transcripts Per Kilobase Million) levels were extracted from GTEx Portal V7 (mean ± SD). **(C)** FLAG-tagged MCART1 localizes to mitochondria. Wild-type HeLa cells transiently expressing FLAG-MCART1 were processed for immunofluorescence detection of the FLAG epitope (cyan) and the mitochondrial inner membrane marker cytochrome c oxidase 4 (COX4) (magenta). The merged image shows the overlap of both channels in white. Scale bar is 10 μm. **(D)** Mitochondrial purification by Mito-IP shows endogenous MCART1 in the mitochondrial fraction. Shown are HA-immunoprecipitates and cell lysates from wild-type cells expressing an HA-mito tag or a control MYC-mito tag. IPs were validated by immunoblotting for the following proteins: CS – citrate synthase; VDAC1 – voltage-dependent anion channel; CALR – calreticulin; GOLGA1 – Golgin subfamily A member; LAMP2 – lysosome-associated membrane glycoprotein; CAT – catalase; RPS6KB1 – Ribosomal protein S6 kinase beta-1. **(E)** Next generation sequencing confirms homozygous 1 or 25 bp frame-shift deletions in the *MCART1* open reading frame in two single-cell derived clones. **(F)** Immunoblot showing loss of MCART1 in Jurkat single cell-derived clones. Lysates prepared from indicated knockout cells were equalized for total protein amount and analyzed by immunoblotting for the levels of the indicated proteins. **(G)** *MCART1*-null cells have a proliferation defect in full RPMI media (mean ± SD; *n* = 3***; *p* < 0.001). The asterisk denotes a statistically significant different between knockout clones and wild-type clones or clones re-expressing the *MCART1* cDNA, respectively. **(H)** Basal and maximal respiration, proton leak, ATP production and spare respiratory capacity are decreased in *MCART1*-null cells. Oxygen consumption rate was measured by Seahorse Extracellular Flux Analysis (mean ± SD; *n* ≥ 13 technical replicates). Graphs were generated from data in the Seahorse experiment in Fig. 1I using the Seahorse Report Generator. **(I)** FLAG-tagged human MCART homologs localize to mitochondria. Wild-type HeLa cells transiently expressing FLAG-constructs were processed for immunofluorescence detection of the FLAG epitope (green) and the mitochondrial inner membrane marker COX4 (magenta). The merged image shows the overlap of both channels in white. Scale bar is 10 μm. **(J)** Immunoblot of *MCART1*-null cells expressing indicated N-terminally FLAG-tagged MCART cDNA constructs. Lysates prepared from indicated cell lines were equalized for total protein amounts and analyzed by immunoblotting for the FLAG-epitope or the levels of the indicated proteins. *indicates endogenous MCART1. ** indicates FLAG-tagged MCART1. **(K)** Human MCART2 but not MCART6 rescues the mitochondrial respiration defect of cells lacking MCART1. Single-cell-derived knockout Jurkat cells were transduced with an empty vector (EV) or cDNAs of human MCART homologs. Asterisks denote statistically significant differences in proliferation in media containing galactose as the carbon source and between the cells expressing the empty vector and the solute carrier homologs. Mean ± SD; *n* = 3; \*\*\*\**P* < 0.0001). **(L)** Within one experiment, the oxygen consumption rate (OCR) and proton efflux rate (PER) were measured by Seahorse extracellular flux analysis with sequential treatment of oligomycin and antimycin A/rotenone to calculate ATP production from glycolysis versus oxidative phosphorylation (mean ± SD; *n* = 5 technical replicates). **(M)** Cellular ATP concentrations in wild-type and *MCART1-null* cells. Data from two independent experiments were combined (mean ± SD; *n* =6). **(N)** Supplementation of RPMI media of cells with metabolites known to alleviate mitochondrial dysfunction does not rescue the proliferation defect of *MCART1*-null cells (mean ± SD; *n* = 3; \*\**P* < 0.01, \*\*\*\**P* < 0.0001). Pyr - pyruvate; HT - hypoxanthine-thymidine. **(O)** Supplementation of RPMI media of cells stably expressing the plasma membrane aspartate transporter SLC1A3 with 10 mM aspartate does not rescue the proliferation defect of *MCART1*-null cells (mean ± SD; *n* = 3; \*\*\**P* < 0.001, \*\*\*\**P* < 0.0001). **(P)** HeLa, HEK-293T and 143B cells lacking MCART1 are deficient in mitochondrial respiration. Proliferation in media containing galactose as the carbon-source was assessed after 4 days. Multiple knockout clones for HeLa and HEK-293T cells are shown, one knockout clone is shown for 143B cells. The bottom panel show loss of MCART1 in knockout clones by western blot.

**Figure S2.**
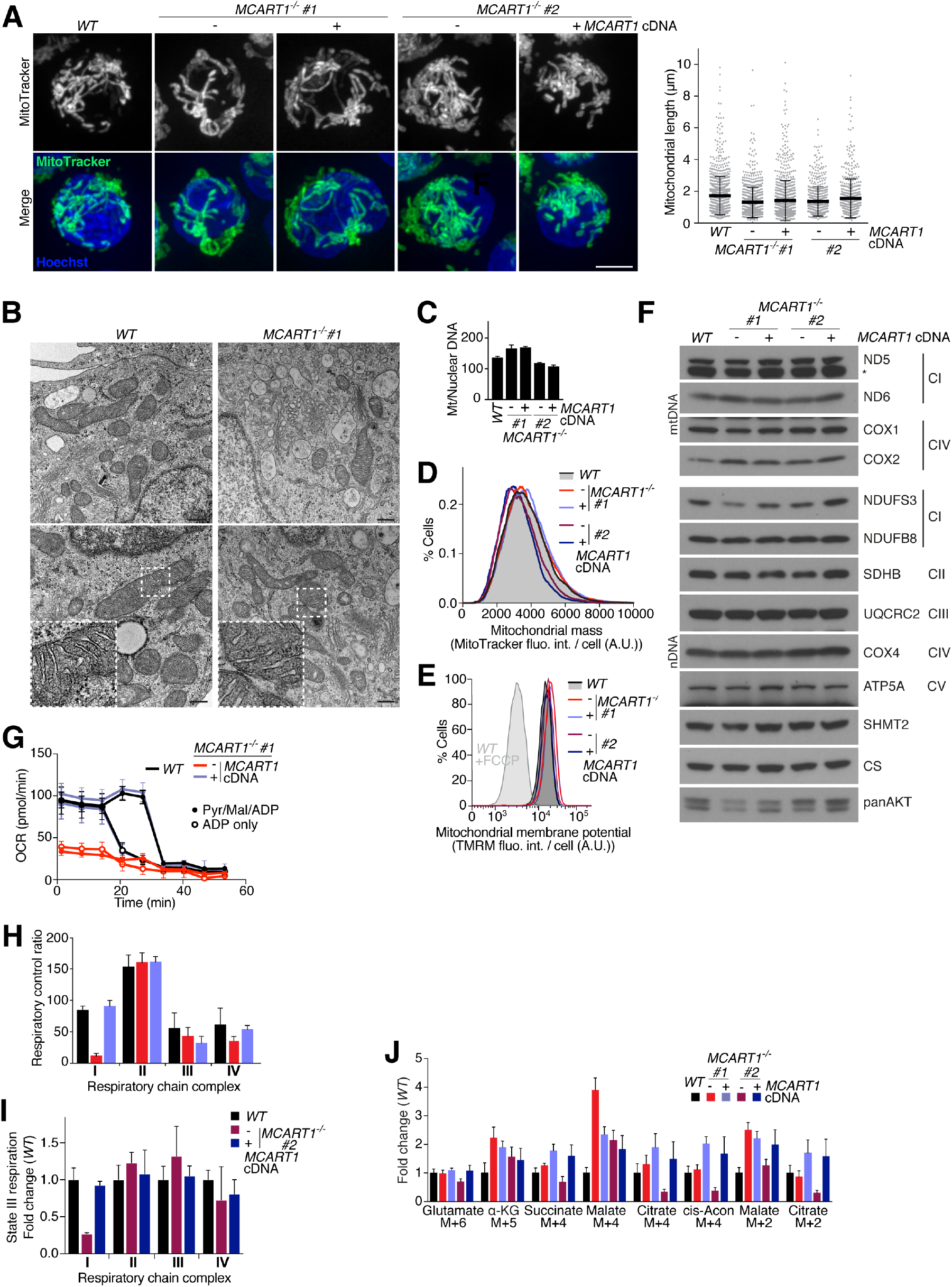
**(A)** Loss of MCART1 does not affect mitochondrial morphology and length. Max intensity z-projections of confocal images of mitochondria visualized by MitoTracker Green (green in merged images) were used to measure mitochondrial length of indicated Jurkat cells. Nuclei were stained with Hoechst DNA stain (blue) (mean ± SD; *n* > 500; \*\*\*\**P* < 0.0001). N.s. – not significant. Scale bar is 5 μm. **(B)** Electron microscopy reveals normal cristae morphology in *MCART1*-null mitochondria. Two 3x magnified inset are shown. Scale bar is 200 nm. **(C)** Loss of MCART1 does not affect mtDNA content (mean ± SD; *n* = 3 technical replicates). Mitochondrial DNA was quantified by qPCR and normalized to genomic DNA. **(D)** Loss of MCART1 does not affect mitochondrial mass per cell as determined by flow cytometry analysis of indicated Jurkat cells stained with MitoTracker Green. The histograms were normalized and smoothened (A.U. – arbitrary units). **(E)** Loss of MCART1 does not affect relative mitochondrial membrane potential as assessed by flow cytometry analysis of Jurkat cells stained with tetramethylrhodamine, methyl ester, and perchlorate (TMRM). Indicated cells were treated with 10 μM FCCP. The histograms were normalized and smoothened. **(F)** Loss of MCART1 only marginally affects protein levels of mitochondrially encoded (upper panel) and nuclear encoded (lower panel) mitochondrial respiratory chain complex subunits. Lysates prepared from indicated cells were equalized for total protein amounts and analyzed by immunoblotting for indicated proteins. CI – complex I; CII – complex II; CIII – complex III; CIV – complex IV; mtDNA – mitochondrially encoded; nDNA – nuclear encoded; CS – citrate synthase. **(G)** *MCART1*-null cells are unable to oxidize exogenous substrate via respiratory complex I. Oxygen consumption rate (OCR) measured by Seahorse extracellular flux analysis of indicated cells permeabilized and supplemented with ADP and complex I substrates or ADP only (mean ± SD; *n* = 3 technical replicates). Mal – malate; perm – permeabilizer; pyr – pyruvate; rot – rotenone. **(H)** Respiratory control ratio determined by Seahorse extracellular flux analysis of indicated cells permeabilized and supplemented with ADP and different respiratory complex substrates (mean ± SD; *n* = 3 technical replicates). **(I)** Complex I-dependent state 3 respiration is diminished in *MCART1*-null clone #2 as determined by Seahorse extracellular flux analysis (mean ± SD; *n* = 3 technical replicates). **(J)** TCA cycle intermediates are still produced in *MCART1*-null cells at the whole cell level. Jurkat cells were incubated in RPMI media containing 2 mM ^13^C_5_,^15^N_2_-glutamine as the sole glutamine source for 2 hours before metabolites were extracted.

**Figure S3.**
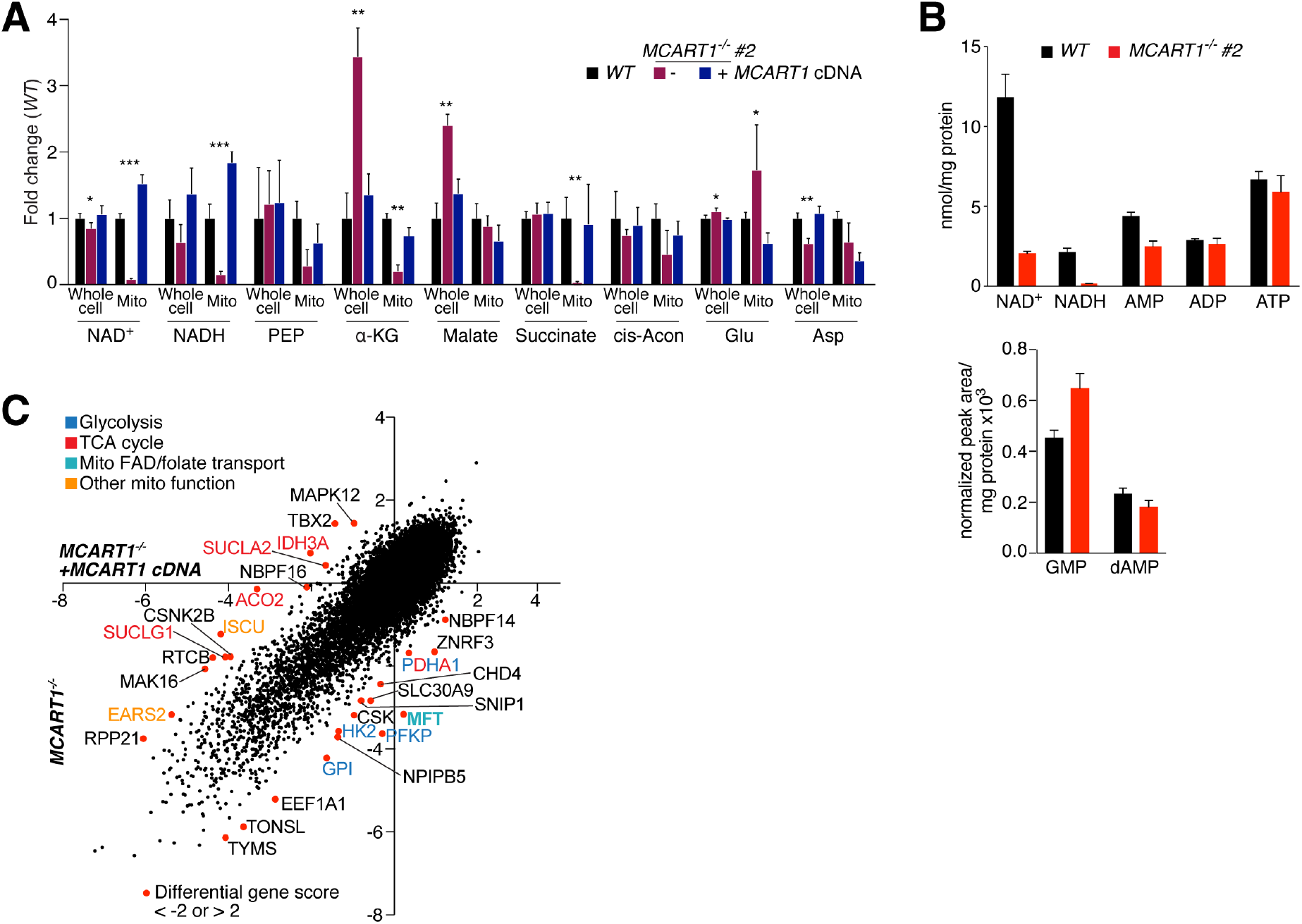
**(A)** NAD^+^ and NADH are depleted in mitochondria of *MCART1-null* clone #2 and TCA cycle intermediates are reduced. Whole cell and mitochondrial (mito) metabolite levels in indicated cells were measured by LC-MS using the Mito-IP method (mean ± SD; *n* = 4). Asterisks denote statistically significant differences of *MCART1-null* samples with both wild-type cells and cells re-expressing the *MCART1* cDNA (\**P* < 0.05, \*\**P* < 0.01, \*\*\**P* < 0.001; PEP - phosphoenolpyruvate; α-KG - α-Ketoglutarate; cis-Acon - cis-aconitate; Glu - glutamate; Asp - aspartate). **(B)** Absolute levels of metabolites in mitochondria isolated by differential centrifugation. Metabolites were measured by LC-MS, quantified based on standard curves and normalized to mitochondrial protein content. Normalized peak areas are shown for GMP and dAMP (mean ± SD; *n* = 3 technical replicates). **(C)** Gene scores from *MCART1*-reexpressing control cells were plotted against those from *MCART1-null* cells. Genes with a differential score of <-2 or >2 are annotated as hits. PDHA1 is labeled in red and blue as it connects glycolysis with the TCA cycle. Mito - mitochondrial; TCA - tricarboxylic acid; FAD - flavin adenine dinucleotide.

**Figure S4.**
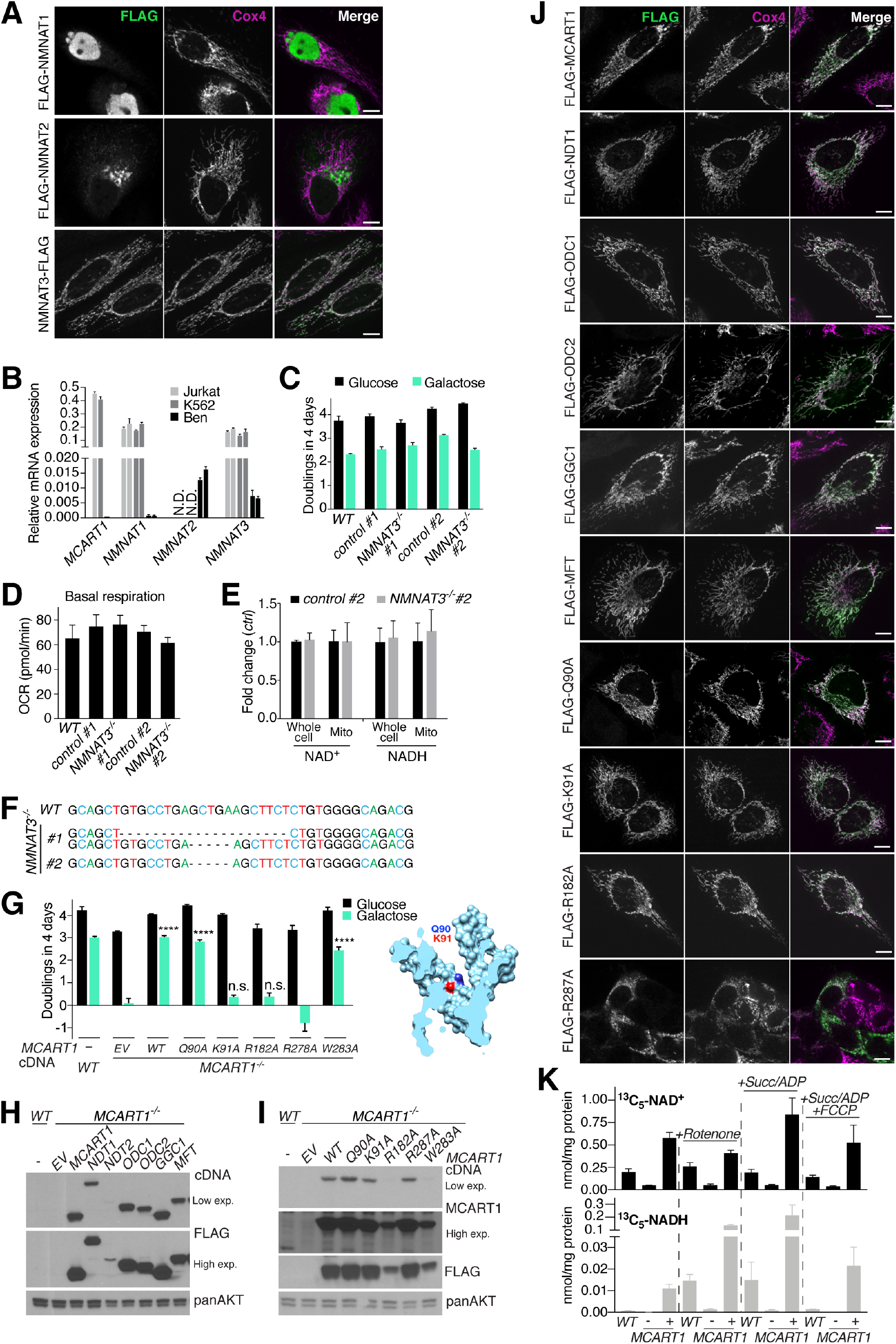
**(A)** FLAG-tagged nicotinamide mononucleotide adenyltransferase (NMNAT) isoforms localize to the nucleus, the Golgi and mitochondria, respectively. Wild-type HeLa cells transiently expressing FLAG-constructs were processed for immunofluorescence detection of the FLAG epitope (green) and the mitochondrial inner membrane marker COX4 (magenta). The merged image shows the overlap of both channels in white. Scale bar is 10 μm. **(B)** Jurkat and K562 cells express *MCART1, NMNAT1*, and *NMNAT3*. mRNA levels were quantified by qPCR relative to *β-ACTIN*. Two primer pairs each were used for NMNAT 1,2 and 3. N.D. – not detected. (Mean ± SD; *n* = 3 technical replicates). **(C)** Human *NMNAT3*-null cells have no mitochondrial respiration defect. Single-cell-derived *NMNAT3* knockout or control Jurkat cells were cultured in media containing glucose or galactose as the carbon source. (Mean ± SD; *n* = 3). N.s. – not significant. **(D)** Human *NMNAT3*-null cells have no mitochondrial respiration defect. A mitochondrial stress test was performed by Seahorse extracellular flux analysis on single-cell-derived *NMNAT3* knockout or control Jurkat cells. (Mean ± SD; *n* > 10 technical replicates). **(E)** Mitochondrial NAD levels do not depend on NMNAT3. (Mean ± SD; *n* = 4). N.s. – not significant. **(F)** Next generation sequencing confirms 20 bp and 5 bp or homozygous 5 bp frame-shift deletions in the *NMNAT3* open reading frame in two single-cell derived clones. **(G)** Mutation of lysine 91 or arginine 278 to alanine abolishes the ability of MCART1 to rescue growth on galactose. *MCART1*-null cells infected with wild-type MCART1 cDNA serve as control cells. (mean ± SD; *n* = 3; \*\*\*\**P* < 0.0001). EV – empty vector. Position of glutamine 90 and lysine 91 in the predicted structure of MCART1 are shown on the right. **(H)** Immunoblot of *MCART1*-null cells expressing indicated N-terminally FLAG-tagged MCART1, MFT or yeast transporter cDNA constructs. Lysates prepared from indicated cells were equalized for total protein amounts and analyzed by immunoblotting for the FLAG-epitope or the levels of the indicated proteins. **(I)** Immunoblot of *MCART1*-null cells expressing indicated N-terminally FLAG-tagged wild-type or K91A mutant MCART1 cDNA constructs. Lysates prepared from indicated cells were equalized for total protein amounts and analyzed by immunoblotting for the FLAG-epitope or the levels of the indicated proteins. **(J)** FLAG-tagged yeast NAD^+^ transporter NDT1, the 2-oxodicarboxylate transporters ODC1 and ODC2, the GTP/GDP transporter GGC1, human MFT and different MCART1 point mutants localize to mitochondria. NDT2 expression levels were not detectable. Wild-type HeLa cells transiently expressing FLAG-constructs were processed for immunofluorescence detection of the FLAG epitope (green) and the mitochondrial inner membrane marker COX4 (magenta). The merged image shows the overlap of both channels in white. Scale bar is 10 μm. **(K)** Transport of NAD^+^ into mitochondria in depends on MCART1. Mitochondria purified from wild-type, *MCART1-null* or *MCART1-null* cells overexpressing the MCART1 cDNA were incubated with 50 μM stably isotope labeled ^13^C_5_-NAD^+^ for 10 min, where indicated in the presence of 5 μM rotenone, succinate and ADP or succinate, ADP and 5 μM FCCP. Levels of taken up ^13^C_5_-NAD^+^ and generated ^13^C_5_-NADH were quantified by LC-MS (mean ± SD; *n* = 3 uptake reactions). The data are representative of three independent experiments.

## Materials and Methods

### Reagents

Reagents were obtained from the following sources: the antibodies that recognize SHMT2 (HPA020549) from Atlas Antibodies; AKT (4691), CALR (12238), Catalase (12980), Citrate Synthase (14309), Cytochrome c oxidase subunit 4 isoform 1 (COX4; 4850), GOLGA1 (13192), RPSS6KB1 (2708), VDAC (4661), the myc (2278) and HA epitopes (3724) and HRP-coupled anti-rabbit secondary antibody as well as Normal Donkey Serum from Cell Signaling Technology (CST); the FLAG epitope from CST (2368) and Sigma (F1804); LAMP2 (sc-18822), TOM20 (sc-11415) and HRP-labeled anti-mouse secondary from Santa Cruz Biotechnology (SCBT); MCART1/SLC25A51 (CSB-PA875649LA01HU) from Cusabio; total OXPHOS Rodent WB Antibody Cocktail (ab110413), ND6 (ab81212) and NDUFS3 (ab177471) from Abcam; Cytochrome c oxidase subunit 1 (COX1; 459600) from Invitrogen; Cytochrome c oxidase subunit 2 (COX2, MTCO2; A-6404) from Life Technologies and the ND1 (55410-1-AP) and ND5 (19703-1-AP) antibodies from Proteintech Group Inc. Antibodies against mitochondrially encoded proteins were validated using ρ^0^-cells. Amino acids, galactose, oligomycin, FCCP, rotenone, antimycin, sodium azide, pyruvic acid, malic acid, ascorbate, adenosine diphosphate, *N,N,N’,N’*-Tetramethyl-*p*-phenylenediamine, malonic acid and succinic acid were from Sigma Aldrich; duroquinol from TCI America; X-tremeGENE 9 and Complete Protease Cocktail from Roche; Alexa 488, 568, and 642-conjugated secondary antibodies and from Invitrogen; anti-HA magnetic beads from ThermoFisher Scientific; glucose from Westnet Inc. (# BM-675); ANTI-FLAG M2 Agarose Affinity Gel and sodium formate from Sigma; egg phosphatidylcholine, *E. coli* total lipids, and the lipid extruder from Avanti Polar Lipids; Bio-Beads SM-2 Adsorbents from Biorad Laboratories; filter membranes for extrusion and supports from Whatman; Cell-Tak from Corning.

### Cell lines and plasmids

The pMXs-IRES-Bsd vector was from Cell Biolabs. The identities of the Jurkat, K562, and HeLa cells used in this study were authenticated by STR profiling. Jurkat cells were used for all functional studies in cells. Sequences of human MCART1, MCART2 and MCART6, MFT and *S. cerevisiae* NDT1 (YIL006W), NDT2 (YEL006W), ODC1 (YPL134C), ODC2 (YOR222W) and GGC1 (YDL198C) were synonymously mutated to remove the proto-spacer adjacent motif (PAM) sequence and/or codon-optimized for expression in human cells. Yeast strains were a gift from Dr. Marina Vai. The wild-type yeast strain derived from <u>CEN.PK</u> 113-7D (*MATa MAL2-8c SUC2*)(*44*). In the *ndt1Δndt2Δ* strain (*36*), NDT1 and NDT2 genes were replaced with the KanMX3 and HphMX4 cassettes by homologous recombination, conferring resistance to G418 and hygromycin B, respectively. For yeast complementation experiments, MCART1 and MCART1K91A sequences were codon-optimized for expression in *S. cerevisiae* and cloned into a centromeric expression plasmid with 5’ and 3’ UTRs of NDT1.

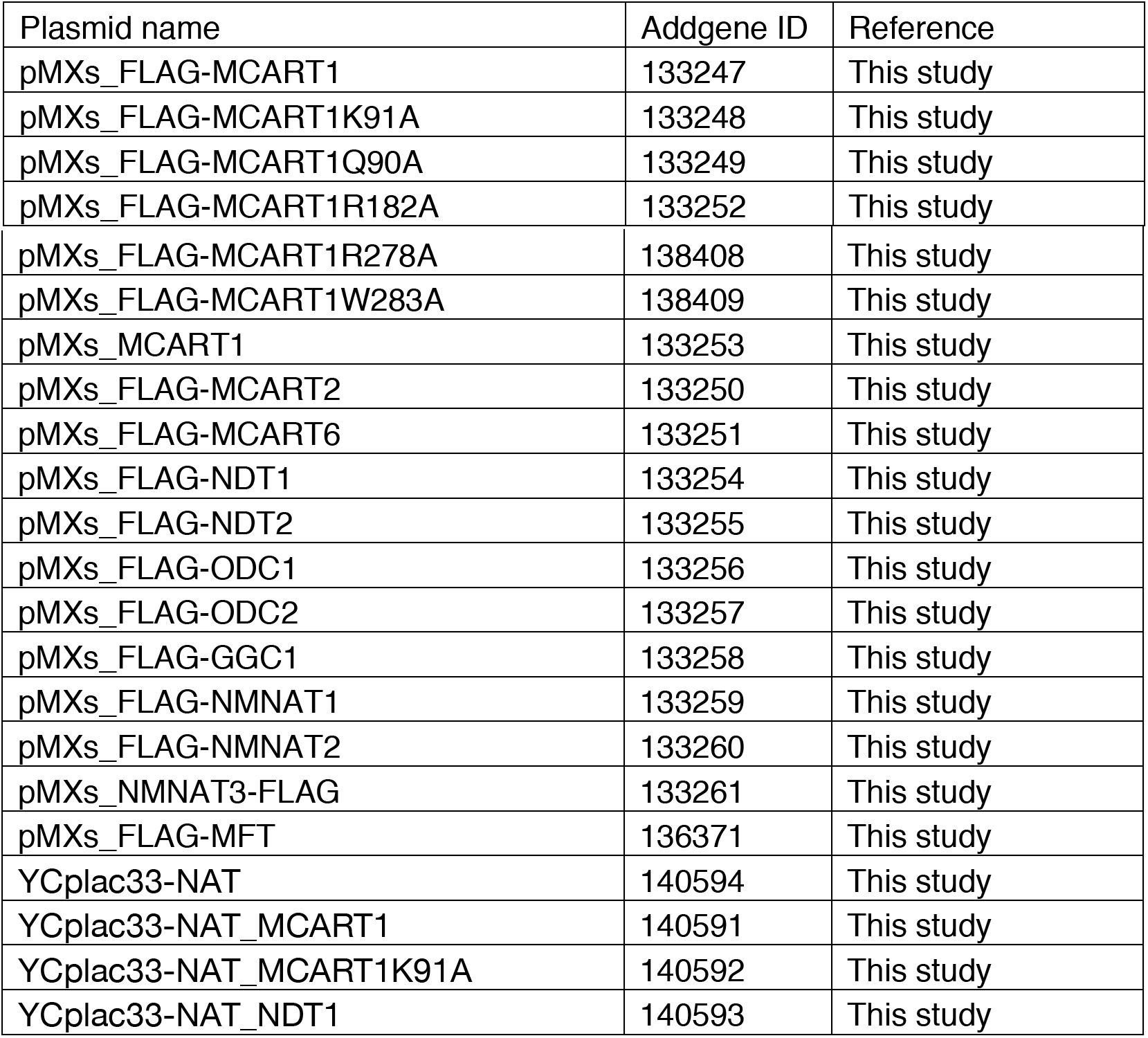

### Cell culture

Unless otherwise indicated, Jurkat and K562 cells were cultured in RPMI (Life Technologies) supplemented with 10% Inactivated Fetal Calf Serum (IFS, Sigma and Gemini), 2 mM glutamine, and penicillin/streptomycin. HeLa, HEK-293T, 14-3B and Ben cells were cultured in DMEM (Life Technologies) supplemented with 10% IFS and penicillin/streptomycin. HEK-293T cells used for virus production were cultured in IMDM (Life Technologies) supplemented with 20% IFS, and penicillin/streptomycin. To compare proliferation of cells in glucose to proliferation in galactose, RPMI without glucose (Life Technologies) was supplemented with dialyzed IFS and either 10 mM glucose or galactose. All cell lines were maintained at 37°C and 5% CO_2_.

### Virus production

HEK-293T cells were co-transfected with the pLentiCRISPR sgRNA library, the VSV-G envelope plasmid and the ΔVPR lentiviral packaging plasmid, or with pMXS plasmids and retroviral packaging plasmids Gag-Pol and VSV-G, using X-TremeGene 9 Transfection Reagent. The culture medium was exchanged 24 hours after transfection with the same medium instead supplemented with 30% IFS. The virus-containing supernatant was collected 48 hours after transfection and spun for 5 min at 400 x g to eliminate cells.

### Transduction of cell lines

Cells were seeded at a density of 1 x10^6^ cells/mL in RPMI containing 8 μg/mL polybrene (EMD Millipore), and then transduced with lentivirus by centrifugation at 2,200 RPM for 45 min at 37°C. After an 18-hour incubation, cells were pelleted to remove virus, washed twice in PBS and then re-seeded into fresh culture medium containing puromycin or blasticidin, and selected for 72 hours.

### CRISPR/Cas9-mediated generation of knockout cell lines

Human MCART1 and NMNAT3 were depleted using the pX330 system and the following sense (S) and antisense (AS) oligonucleotides:

sgMCART1_3 (S): caccgGAGATGAAGCATTACTTGTG
sgMCART1_3 (AS): aaacCACAAGTAATGCTTCATCTCc
sgNMNAT3_9 (S): caccgCCACAGAGAAGCTTCAGCTC
sgNMNAT3_9 (AS): aaacGAGCTGAAGCTTCTCTGTGGc

1 million Jurkat cells were electroporated with the 2.5 μg of sgRNA plasmid and GFP control plasmid at a 10:1 ratio using an Amaxa Cell Line Nucleofector Kit V and an Amaxa™ Nucleofector™ II (Lonza). HeLa, 143B and HEK-293T cells were transfected using Fugene transfection reagent. GFP-positive cells were single-cell FACS-sorted into 96-well plates containing media supplemented with 30% IFS and 50% conditioned media. Cell clones with MCART1 knockouts were identified by western blotting and confirmed by next generation sequencing at the MGH CCIB DNA core. Two different knockout clones of Jurkat cells (designated #1 and #2) were used for experiments. Between experiments *MCART1-null* cells were maintained in media supplemented with 30% IFS, 2 mM L-glutamine, 1 mM uridine and 0.1 mM/16 μM hypoxanthine/thymidine. NMNAT3 knockout clones were identified by next generation sequencing.

### CRISPR-Cas9 synthetic lethality negative selection genetic screen

*MCART1*-null or control cells re-expressing the guide-resistant *MCART1* cDNA were transduced with a genome-wide sgRNA library (*7, 45*). 48 hours after infection, cells were selected with puromycin for 72 hours. Subsequently, cells were passaged every other day at a seeding density of 250,000 cells/ml until reaching ~14 population doublings (PDs). DNA was extracted from 80 x10^6^ cells using the QIAamp DNA Blood Maxi Kit (QIAGEN). sgRNA inserts were PCR amplified using ExTaq DNA Polymerase (Takara). The resultant PCR products were purified and sequenced on a HiSeq 2500 (Illumina) (primer sequences provided below) to monitor the change in the abundance of each sgRNA between the initial and final cell populations.

Primer sequences for sgRNA quantification

Forward:

AATGATACGGCGACCACCGAGATCTACACGAATACTGCCATTTGTCTCAAGATCTA Reverse: CAAGCAGAAGACGGCATACGAGATCnnnnnnTTTCTTGGGTAGTTTGCAGTTTT (nnnnnn denotes the sample barcode).

Illumina sequencing primer

CGGTGCCACTTTTTCAAGTTGATAACGGACTAGCCTTATTTTAACTTGCTATTTCTAG CTCTAAAAC

Illumina indexing primer

TTTCAAGTTACGGTAAGCATATGATAGTCCATTTTAAAACATAATTTTAAAACTGCAA ACTACCCAAGAAA

Sequencing reads were aligned to the sgRNA library and the abundance of each sgRNA was calculated. sgRNAs with less than 50 counts in the initial cell pool were removed from downstream analyses. The log2fold-change in abundance of each sgRNA was calculated for each treatment condition after adding a pseudocount of one. A pseudocount of one was added to all sgRNAs and counts were normalized by number of reads in each sample multiplied by one million. Gene scores were defined as the average log2fold-change in the abundance of all sgRNAs targeting a given gene between the initial and final cell populations and calculated for both cell lines and Z-score normalized. The differential gene score was calculated as the difference in gene scores between cell lines.

### Cell proliferation assays

10,000 cells per well were seeded into 96-well plates in triplicate. Cell titer glo reagent (Promega) was added to one plate 1 hour after seeding and luminescence was measured, while a second plate was read-out 4 days after seeding. Number of doublings in 4 days was determined by calculating the log2 fold-change in signal between day 0 and 4. For cell counting experiments, 100,000 cells/ml were seeded in a 6-well plate in triplicates and counted immediately after seeding and from then on every 24 hours for 5 sequential days. For metabolite rescue experiments, 1 mM pyruvate and 100 μg/ml uridine, 1 mM formate or 0.1 mM/16 μM hypoxanthine/thymidine were added at the beginning of the culture period. For proliferation assays with HeLa, HEK-293T or 143B cells, 2,000 cells per well were seeded and cultured in media containing 30% IFS (and glutamine, uridine and hypoxanthine/thymidine) supplemented with 11 mM glucose or galactose for three days before switching to media of the respective carbon source supplemented with 10% dialyzed IFS only for another 24 hours before readout.

### mRNA quantification by qPCR

RNA was extracted from cells using the RNeasy kit according to the manufacturer’s instructions (Qiagen) and RNA reverse transcribed using the SuperScript III Reverse Transcriptase (Thermo Fisher Scientific). The following primers were used to assess mRNA levels of *MCART1, NMNAT1, NMNAT2 and NMNAT3* by qPCR by normalizing Ct values to those of *β-ACTIN*. MCART1_L: TAAGGAGCATCTGCCTACCG; MCART1_R: CCCAACATGGCACCCAATAG; NMNAT1_L1: AAGCTGTGCCAAAGGTCAAG; NMNAT1_R1: TTCCAGCCCGAGTAACACAT; NMNAT1_L2: GTGGTTCTCCTTGCTTGTGG; NMNAT1_R2: TAGTCCTTGGCCAGCTCAAA; NMNAT2_L1: GTGGAGCGTTTCACCTTTGT; NMNAT2_R1: CACCTCCATATCTGCCTCGT; NMNAT2_L2: CCGTCTCATCATGTGTCAGC; NMNAT2_R2: AGGTGTCATGGAAGGTGTGT; NMNAT3_L1: GATGCGCACATCCAGGAAAT; NMNAT3_R1: TTGGCCAGGTGAATGTTGTG; NMNAT3_L2: ATGGGAAGAAAGACCTCGCA; NMNAT3_R2: CCTCAGCACCTTCACTGTCT; β-ACTIN_L; AGGATGGCAAGGGACTTCCTG; β-ACTIN_R: AATGTGGCCGAGGACTTTGAT.

### Immunofluorescence assays and STED imaging

For immunofluorescence assays 50,000 HeLa cells were plated in a 24-well glass bottom imaging plate (Cellvis, Mountain View, CA) and transfected with 500 ng of the cDNAs for FLAG constructs 16 hours later. 48 hours after transfection, cells were rinsed twice with PBS and fixed with 3% paraformaldehyde with 0.1% glutaraldehyde in PBS for 10 minutes. The fixation and all subsequent steps were performed at room temperature. Cells were rinsed three times with PBS and permeabilized with 0.3% NP40, 0.05% Triton X-100, 0.1% BSA in PBS for 3 minutes. After rinsing three times with wash buffer (0.05% NP40, 0.05% Triton-X 100, 0.2% BSA in PBS) samples were blocked for 1 hour in blocking buffer (0.05% NP40, 0.05% Triton-X 100, 5% Normal Donkey Serum). The samples were incubated with primary antibody in blocking buffer for 1 hour, washed three times with wash buffer, and incubated with secondary antibodies produced in donkey (diluted 1:500 in blocking buffer) for 30 minutes in the dark, washed three times with wash buffer, and rinsed three times with PBS. The primary antibodies used were directed against COX4 (CST; 1:250 dilution), the FLAG epitope (Sigma, 1:500 dilution) and TOM20 (SCBT, 1:500). Secondaries antibodies conjugated with Alexa 488 and 568 were used for confocal microscopy. Images were acquired on a Zeiss AxioVert200M microscope with a 63X oil immersion objective and a Yokogawa CSU-22 spinning disk confocal head with a Borealis modification (Spectral Applied Research/Andor) and a Hamamatsu ORCA-ER CCD camera. The MetaMorph software package (Molecular Devices) was used to control the hardware and image acquisition. The excitation lasers used to capture the images were 488 nm and 561 nm. Images were processed with FIJI (*46*). STED imaging was carried out on a Leica TCS SP8 STED 3X setup with an HC PL APO 100x/1.40 oil STEDwhite objective. Samples were fixed as described above. FLAG was detected using Alexa 594 secondary antibodies, COX4 with Atto647N (Sigma Aldrich), and Tom20 with Alexa 488. 660nm and 775nm or 592nm depletion lasers were used. Images were deconvolved using the Adaptive Lightning strategy (Leica). Line profiles were generated from the raw data using FIJI.

### MS-based metabolomics and quantification of metabolite abundances

Metabolite abundance using LC/MS-based metabolomics was measured and quantified as previously described (*11*). Briefly, Jurkat cells were seeded at a density of 0.6 x10^6^ per ml. 24 hours later, 1.5-2 x10^6^ cells were harvested, washed once in ice-cold 0.9% saline prepared with LC-MS-grade water, and extracted with 80% methanol containing 500 nM isotope-labeled amino acids as internal standards (Cambridge Isotope Laboratories). Where indicated in the section “Mitochondrial isolations for metabolite analyses” a different extraction method was used to specifically quantify NAD levels. The samples were vortexed for 10 min at 4°C and centrifuged at 17,000 x g. The supernatant was dried by vacuum centrifugation at 4°C. Samples were stored at −80°C until analyzed. On the day of analysis, samples were resuspended in 50-100 μL of LC-MS-grade water and the insoluble fraction was cleared by centrifugation at 15,000 rpm. The supernatant was then analyzed as previously described by LC-MS (*11, 21*). Amino acids were normalized to their respective internal standards, TCA cycle intermediates, malate, carnitine and C5-carnitine were normalized to the glutamate internal standard. NAD^+^ and NADH were normalized to the valine internal standard.

### Glucose consumption and metabolite secretion assay

For media metabolite extraction, 6 wells of a 6-well plate were seeded with 300,000 Jurkat cells in 2 ml RPMI. The next day, cells were washed in PBS and resuspended in media. 1 ml of the upper three wells was collected and centrifuged at 3000 RPM for 5 min. to pellet cells. For the remaining media, the number of cells was counted. Two days later, the same procedure was repeated for the lower three wells. Metabolites were extracted from the media with a 75/25/0.2 extraction mix acetonitrile/methanol/formic acid with internal standards by vortexing for 10 min at 4°C. Samples were centrifuged at 17,000 x g and the supernatant was analyzed by LC/MS. To calculate metabolite secretion or consumption rates, the difference in concentration between day 2 and day 0 were divided by the area under the growth curve according to (*47*).

### Mitochondrial isolations for immunoblot analyses

30 x10^6^ Jurkat cells expressing the HA-mito tag or a control tag were washed 1x in PBS, 1x in KPBS according to (*21*). 5 μl of the cell suspension in 1 ml KPBS was lysed in 50 μl of 1% Triton lysis buffer to obtain whole cell protein levels. The rest was lysed using 8 strokes with a 30 ½ G needle. Lysates were spun for 1 min at 1000 x g to pellet unbroken cells, and subsequently incubated with 100 μl HA-magnetic beads for 4 min. Beads were washed 3x in KPBS, and mitochondria lysed in 50 μl lysis buffer for 10 min. Beads were removed using the magnet, and samples were spun 10 min at 17,000 x g to remove residual beads and insoluble material. SDS-PAGE loading dye was added to each sample, and 6 μl of whole cell lysate and 9 μl of the mitochondrial fraction were analyzed by SDS-PAGE.

### Mitochondrial isolations for metabolite analyses

Mitochondria were isolated using the Mito-IP method as described above for immunoblotting, except cells were disrupted with 20 strokes in a homogenizer containing a pure PTFE head (VWR International) and two strokes with a dounce tissue grinder with tight-fitting pestle (DWK Life Sciences Kimble Kontes). 850 μl of the final suspension of beads with bound mitochondria in 1 ml KPBS were used for metabolite extraction and the remaining 150 μl for immunoblotting to determine mitochondrial capture efficiency. Metabolites were extracted with 50 μl 80% methanol containing internal standards. To obtain paired whole cell metabolite quantification, 25 μl of the initial cell suspension in 1 ml KPBS were extracted in 225 μl of 80% methanol with internal standards. In experiments in Fig. 4 and figs. S3 and S4 an extraction buffer consisting of 2:2:1 acetonitrile:methanol:water was used to preserve NAD^+^ and NADH followed by quenching with 15% (w/v) ammonium bicarbonate (8.7μl per 100μl solvent; adapted from (*22*)). 5 μl of the mitochondrial extract was injected for mass spectrometry analysis. Mitochondrial metabolite levels were normalized based on the citrate synthase signal determined by western blot. For the experiment in fig, S3B, mitochondria were isolated as described in mitochondrial uptake assays and washed 1x in KPBS before metabolite extraction. Separate replicates were used for protein quantification by the BCA method.

### Mitochondrial transport assays

Mitochondria were isolated as described previously (*48*). Cells (at least ~1 g starting material) grown in shaking culture (in glucose-containing RPMI as described above, with 0.1% pluronic (Gibco)) were resuspended in low tonicity buffer (100 mM sucrose, 10 mM MOPS pH 7.2, 1 mM EGTA, 0.1% BSA) and disrupted with 20 strokes in a homogenizer containing a pure PTFE head (VWR International) and two strokes with a dounce tissue grinder with tight-fitting pestle (DWK Life Sciences Kimble Kontes). After adjusting tonicity, lysates were spun twice at 950 x g to remove unbroken cells and nuclei, and mitochondria were pelleted by centrifuging at 10,000 x g for 10 min. The supernatant as well as the top, white part of the pellet containing other membranes was aspirated and the brown/yellow mitochondrial pellet was washed in isolation buffer (210 mM mannitol, 70 mM sucrose, 10 mM MOPS pH 7.2, 1 mM EGTA, 0.1% BSA) before resuspending in mitochondria incubation media (125 mM KCl, 10 mM MOPS pH 7.2, 2 mM MgCl_2_, 2 mM KH_2_PO_4_, 10 mM NaCl, 1 mM EGTA). Mitochondrial preparations were adjusted for protein content. 100 μl volume uptake reactions with 50 μM ^13^C_5_-NAD^+^ (Cambridge Isotope Laboratories) were initiated by adding substrate to 80 μg of mitochondria in incubation media containing 0.1% BSA and 100 μM unlabeled NAD^+^ at 30°C. 5 μM rotenone, 10 mM succinate/1 μM ADP and 5 μM FCCP were added where indicated. For competition experiments, 500 μM labeled NAD^+^ was added to mitochondria preincubated with 50 μM unlabeled NAD^+^. Additionally, unlabeled NAD^+^, NADH or NMN were added at 2 or 5 mM where indicated. After 10 min 700 μl ice-cold KPBS were added to the reaction and the samples were collected by centrifugation at 10,000 x g for 3 min at 4°C. Mitochondrial pellets were washed once in ice-cold KPBS before extracting the metabolites in 160μl extraction buffer (2:2:1 acetonitrile:methanol:water followed by quenching with 12μl 15% (w/v) ammonium bicarbonate). Absolute levels of labeled NAD^+^ and NADH were quantified based on a standard curve. ^15^N-^13^C_5_-Valine was used as an internal standard.

### Yeast complementation experiments

*Ndt1Δndt2Δ* cells were transformed with empty vector or plasmids containing *S. cerevisiae NDT1*, human *MCART1* or the K91A point mutant with 5’ and 3’ UTRs of *NDT1* and plated onto standard YPD plates with 100 μg/ml Nourseothricin (NAT). For growth and metabolomics experiments yeast were cultured in baffled flasks shaking at 250 rpm at 30°C in synthetic minimal media consisting of yeast nitrogen base without amino acids, ammonium sulfate and 2% ethanol as a carbon source with 100 μg/ml NAT. For growth assays, cultures were inoculated at an OD_600_ of 0.1 from pre-cultures. Growth curves were fitted based on the average of the two independent clones except for wild-type cells. Mitochondria were isolated according to a published protocol (*49*) and extracted as described above.

### Glutamine tracing experiments

For glutamine tracing experiments in whole cells, cells were incubated in RPMI containing 2 mM ^13^C_5_,^15^N_2_-glutamine (Cambridge Isotope Labs) as the sole glutamine source for 2 hours before metabolites were extracted and quantified as described above with the exception that no internal standards were added to the extraction buffer. To measure TCA cycle flux in isolated mitochondria, mitochondria were purified by Mito-IP as described above. Mitochondria bound to magnetic beads were incubated in 100 μl KPBS containing 4 mM ^13^C_5_,^15^N_2_-glutamine, 0.5 mM malate, 1 mM ADP at 33°C for 2.3 hours with rocking. The reaction was stopped and metabolites extracted by addition of 150 μl ice-cold acetonitrile. Samples were vortexed and centrifuged at 17,000 x g for 10min. 4 μl of a 1:10 dilution in 80% methanol was injected for mass spectrometric detection. Metabolites were normalized based on the citrate synthase signal determined by western blot.

### Seahorse extracellular flux analyses

Oxygen consumption rates (OCR) of intact cells were measured using an XFe96 Extracellular Flux Analyzer (Agilent). 100,000 Jurkat cells were seeded on Seahorse XFe96 culture plates coated with Cell-tak and assayed after incubation at 37°C for 1 h. Three basal OCR measurements were taken, followed by sequential injections of 1 oligomycin, 3 μM FCCP, and 1 μM antimycin A, taking three measurements following each treatment. Cellular respiration was calculated by subtracting the OCR after Antimycin A treatment from the basal or FCCP-stimulated OCR. ATP synthesis was measured with the Seahorse XF Real-Time ATP Rate Assay kit (Agilent) according to the manufacturer’s instructions. Activity measurements for each respiratory chain complex in permeabilized cells were performed according to (*50*). Briefly, after seeding of cells the media was changed to Mannitol and sucrose (MAS)-BSA buffer (70 mM sucrose, 220 mM Mannitol, 10 mM KH2PO4, 5 mM MgCl2, 2 mM HEPES pH 7.2, 1 mM EGTA, 0.4% BSA) and flux measurements were started. After three basal measurements, cells were permeabilized by injection of XF Plasma Membrane Permeabilizer (Agilent; 1nM final concentration) together with 1mM ADP and the following respiratory complex substrates or ADP only: complex I - pyruvate/malate (5 mM/2.5 mM); complex II - succinate/rotenone (10mM/1μM); complex III - duroquinol (0.5mM); complex IV - N,N,N,N-tetramethyl-p-phenylenediamine/ascorbate (0.5mM/2mM final concentration). These were followed by injections with Oligomycin (1 μg/ml final), and respective complex inhibitors (complex I - 1μM rotenone, complex II,III - 20 μM antimycin A, complex IV - 20 mM sodium azide). Wells in which cells were lifted off the plate during the assay were excluded from the analysis.

### Characterization of mitochondria

For quantification of mitochondrial mass and morphology, Jurkat cells were stained with MitoTracker Green (Life Technologies, M22426) at 25 nM for 1h before analysis by flow cytometry or fluorescence microscopy. For microscopy, nuclei were stained with Hoechst 33342 fluorescent stain (Molecular Probes) at 2 μg/ml and z-stacks with 250 nm step size were taken at 100x magnification. FIJI was used to generate max intensity z-projections and measure mitochondrial length (*46*). To measure mitochondrial membrane potential cells were stained with 200 nM tetramethylrhodamine, methyl ester, perchlorate (TMRM; Life Technologies, T668) in RPMI for 20 min at 37°C, washed once with PBS, and resuspended in fresh PBS for flow cytometry analysis of live cells. Where indicated, cells were incubated with 10 μM FCCP for 10 min prior to adding TMRM dye. For ultrastructural analysis by electron microscopy, cells were were fixed in 2.5% glutaraldehyde, 3% paraformaldehyde with 5% sucrose in 0.1M sodium cacodylate buffer (pH 7.4), pelletted, and post fixed in 1% OsO4 in veronal-acetate buffer. The cells were stained en block overnight with 0.5% uranyl acetate in veronal-acetate buffer (pH6.0), then dehydrated and embedded in Embed-812 resin. Sections were cut on a Leica EM UC7 ultra microtome with a Diatome diamond knife at a thickness setting of 50 nm, stained with 2% uranyl acetate, and lead citrate. The sections were examined using a FEI Tecnai spirit at 80KV and photographed with an AMT ccd camera. Analysis of mtDNA copy number was performed as previously described (*13*). Briefly, genomic and mitochondrial DNA were extracted from cells using the QIAamp DNA mini kit according to the manufacturer’s instructions (Qiagen). The following primers targeting the mitochondrial gene ND1 and the nuclear gene RUNX2 were used to assess mtDNA copy number by qPCR by normalizing Ct values of ND1 to those of RUNX2. ND1_F: CCC TAA AAC CCG CCA CAT CT; ND1_R: GAG CGA TGG TGA GAG CTA AGG T; RUNX2_F: CGC ATT CCT CAT CCC AGT ATG; RUNX2_R: AAA GGA CTT GGT GCA GAG TTC AG. Jurkat whole cell lysates for immunoblot analysis of mitochondrial (and other) proteins were prepared by lysis in 1% Triton lysis buffer.

### Bioinformatics analyses

MCART1 (Q9H1U9; S2551_HUMAN) topology was predicted using Protter (*10*). The Broad Institute Achilles CRISPR data set (*8*) was analyzed in excel using the in-build correlation function to calculate MCART1 correlation with all genes in the dataset. In addition, the Achilles data was further analyzed in “R” (version 3.3.3, x64) using custom written scripts. The limma package was used to generate barcode plots. Gene Ontology (GO) terms used for barcode plots are ETC: GO:0022900; TCA cycle: GO:0006099; Mitochondrial DNA replication: GO:0006264; Mitophagy: GO:0000423; Mitochondrial fusion: GO:0008053; Mitochondrial transmembrane transport: GO:1990542; Fatty acid beta-oxidation: GO:0006635. All groups were filtered for human genes only. For construction of the phylogenetic tree, protein sequences of all members of the human SLC25 family (obtained from (*51*)) were aligned using MUSCLE (*52*). The PHYLIP proml module (*53*) was used to construct the phylogenetic tree and FigTree software v.1.4.3 to visualize it. Percent sequence identities and similarities were calculated with the NCBI blastp tool. Graphpad Prism 7 software was used to generate the heat map of *MCART* RNA expression based on data from the Cancer Cell Line Encyclopedia (broadinstitute.org/ccle). *MCART* TPM (Transcripts Per Kilobase Million) levels in normal tissues were extracted from GTEx Portal V7. The MCART1 structure was modeled using Memoir membrane protein modelling pipeline (*54*) based on the structure of the bovine mitochondrial ADP/ATP carrier in complex with carboxyatractyloside ((*32*);PDB #1OKC). The high-coverage model was visualized using Chimera (*55*).

### Statistical analyses

Two-tailed t tests were used for comparison between two groups. All comparisons were two-sided, and P values of less than 0.05 were considered to indicate statistical significance. All error bars denote standard deviations between biological replicates unless indicated otherwise. Where fold changes are shown, the average of control or wild-type replicates was normalized to 1 and other samples were normalized accordingly.

## References

1. R. H. Houtkooper, C. Canto, R. J. Wanders, J. Auwerx, The secret life of NAD+: an old metabolite controlling new metabolic signaling pathways. Endocr Rev 31, 194–223 (2010).

2. E. Verdin, NAD(+) in aging, metabolism, and neurodegeneration. Science 350, 1208–1213 (2015).

3. L. Rajman, K. Chwalek, D. A. Sinclair, Therapeutic Potential of NAD-Boosting Molecules: The In Vivo Evidence. Cell Metab 27, 529–547 (2018).

4. L. R. Stein, S. Imai, The dynamic regulation of NAD metabolism in mitochondria. Trends in endocrinology and metabolism: TEM 23, 420–428 (2012).

5. S. Todisco, G. Agrimi, A. Castegna, F. Palmieri, Identification of the mitochondrial NAD+ transporter in Saccharomyces cerevisiae. The Journal of biological chemistry 281, 1524–1531 (2006).

6. F. Palmieri et al., Molecular Identification and Functional Characterization of Arabidopsis thaliana Mitochondrial and Chloroplastic NAD(+) Carrier Proteins. Journal of Biological Chemistry 284, 31249–31259 (2009).

7. T. Wang et al., Gene Essentiality Profiling Reveals Gene Networks and Synthetic Lethal Interactions with Oncogenic Ras. Cell 168, 890–903 e815 (2017).

8. R. M. Meyers et al., Computational correction of copy number effect improves specificity of CRISPR-Cas9 essentiality screens in cancer cells. Nature genetics 49, 1779–1784 (2017).

9. Z.-Y. L. Hana Antonicka, Alexandre Janer, Woranontee Weraarpachai, View ORCID ProfileAnne-Claude Gingras, Eric A. Shoubridge, A high-density human mitochondrial proximity interaction network. bioRxiv preprint, (2020).

10. U. Omasits, C. H. Ahrens, S. Muller, B. Wollscheid, Protter: interactive protein feature visualization and integration with experimental proteomic data. Bioinformatics 30, 884–886 (2014).

11. K. Birsoy, Wang, T., Chen, W.W., Freinkman, E., Abu-Remaileh, M., Sabatini, D.M., An Essential Role of the Mitochondrial Electron Transport Chain in Cell Proliferation Is to Enable Aspartate Synthesis. Cell 162, 540–551 (2015).

12. M. P. King, G. Attardi, Human cells lacking mtDNA: repopulation with exogenous mitochondria by complementation. Science 246, 500–503 (1989).

13. N. Kory et al., SFXN1 is a mitochondrial serine transporter required for one-carbon metabolism. Science 362, eaat9528 (2018).

14. G. S. Ducker et al., Reversal of Cytosolic One-Carbon Flux Compensates for Loss of the Mitochondrial Folate Pathway. Cell Metab 23, 1140–1153 (2016).

15. H. Patel, E. D. Pietro, R. E. MacKenzie, Mammalian fibroblasts lacking mitochondrial NAD+-dependent methylenetetrahydrofolate dehydrogenase-cyclohydrolase are glycine auxotrophs. The Journal of biological chemistry 278, 19436–19441 (2003).

16. X. R. Bao et al., Mitochondrial dysfunction remodels one-carbon metabolism in human cells. Elife 5, (2016).

17. I. Martínez-Reyes et al., TCA cycle and mitochondrial membrane potential are necessary for diverse biological functions. Molecular cell 61, 199–209 (2016).

18. A. R. Grassian et al., IDH1 mutations alter citric acid cycle metabolism and increase dependence on oxidative mitochondrial metabolism. Cancer Res 74, 3317–3331 (2014).

19. C. A. Lewis et al., Tracing compartmentalized NADPH metabolism in the cytosol and mitochondria of mammalian cells. Molecular cell 55, 253–263 (2014).

20. R. P. Hausinger, FeII/alpha-ketoglutarate-dependent hydroxylases and related enzymes. Critical reviews in biochemistry and molecular biology 39, 21–68 (2004).

21. W. W. Chen, E. Freinkman, T. Wang, K. Birsoy, D. M. Sabatini, Absolute Quantification of Matrix Metabolites Reveals the Dynamics of Mitochondrial Metabolism. Cell 166, 1324–1337 e1311 (2016).

22. W. Lu, L. Wang, L. Chen, S. Hui, J. D. Rabinowitz, Extraction and Quantitation of Nicotinamide Adenine Dinucleotide Redox Cofactors. Antioxid Redox Signal 28, 167–179 (2018).

23. D. Hellebrekers et al., Novel SLC25A32 mutation in a patient with a severe neuromuscular phenotype. Eur J Hum Genet 25, 886–888 (2017).

24. M. Schiff et al., SLC25A32 Mutations and Riboflavin-Responsive Exercise Intolerance. The New England journal of medicine 374, 795–797 (2016).

25. E. A. McCarthy, S. A. Titus, S. M. Taylor, C. Jackson-Cook, R. G. Moran, A mutation inactivating the mitochondrial inner membrane folate transporter creates a glycine requirement for survival of chinese hamster cells. The Journal of biological chemistry 279, 33829–33836 (2004).

26. S. A. Titus, R. G. Moran, Retrovirally mediated complementation of the glyB phenotype - Cloning of a human gene encoding the carrier for entry of folates into mitochondria. Journal of Biological Chemistry 275, 36811–36817 (2000).

27. A. N. Spaan et al., Identification of the human mitochondrial FAD transporter and its potential role in multiple acyl-CoA dehydrogenase deficiency. Mol Genet Metab 86, 441–447 (2005).

28. A. Nikiforov, C. Dolle, M. Niere, M. Ziegler, Pathways and subcellular compartmentation of NAD biosynthesis in human cells: from entry of extracellular precursors to mitochondrial NAD generation. The Journal of biological chemistry 286, 21767–21778 (2011).

29. L. Liu et al., Quantitative Analysis of NAD Synthesis-Breakdown Fluxes. Cell Metab 27, 1067–1080 e1065 (2018).

30. M. Barile, S. Passarella, G. Danese, E. Quagliariello, Rat liver mitochondria can synthesize nicotinamide adenine dinucleotide from nicotinamide mononucleotide and ATP via a putative matrix nicotinamide mononucleotide adenylyltransferase. Biochem Mol Biol Int 38, 297–306 (1996).

31. A. Davila et al., Nicotinamide adenine dinucleotide is transported into mammalian mitochondria. Elife 7, (2018).

32. E. Pebay-Peyroula et al., Structure of mitochondrial ADP/ATP carrier in complex with carboxyatractyloside. Nature 426, 39–44 (2003).

33. J. J. Ruprecht et al., Structures of yeast mitochondrial ADP/ATP carriers support a domain-based alternating-access transport mechanism. Proceedings of the National Academy of Sciences of the United States of America 111, E426–434 (2014).

34. J. J. Ruprecht et al., The Molecular Mechanism of Transport by the Mitochondrial ADP/ATP Carrier. Cell 176, 435–447 e415 (2019).

35. D. R. Nelson, J. E. Lawson, M. Klingenberg, M. G. Douglas, Site-directed mutagenesis of the yeast mitochondrial ADP/ATP translocator. Six arginines and one lysine are essential. Journal of molecular biology 230, 1159–1170 (1993).

36. I. Orlandi, G. Stamerra, M. Vai, Altered Expression of Mitochondrial NAD(+) Carriers Influences Yeast Chronological Lifespan by Modulating Cytosolic and Mitochondrial Metabolism. Front Genet 9, 676 (2018).

37. G. Agrimi et al., Deletion or overexpression of mitochondrial NAD+ carriers in Saccharomyces cerevisiae alters cellular NAD and ATP contents and affects mitochondrial metabolism and the rate of glycolysis. Appl Environ Microbiol 77, 2239–2246 (2011).

38. M. R. VanLinden et al., Subcellular Distribution of NAD(+) between Cytosol and Mitochondria Determines the Metabolic Profile of Human Cells. Journal of Biological Chemistry 290, 27644–27659 (2015).

39. X. A. Cambronne et al., Biosensor reveals multiple sources for mitochondrial NAD(+). Science 352, 1474–1477 (2016).

40. M. Yamamoto et al., Nmnat3 Is Dispensable in Mitochondrial NAD Level Maintenance In Vivo. Plos One 11, (2016).

41. B. J. Floyd et al., Mitochondrial Protein Interaction Mapping Identifies Regulators of Respiratory Chain Function. Molecular cell 63, 621–632 (2016).

42. S. I. Imai, L. Guarente, It takes two to tango: NAD(+) and sirtuins in aging/longevity control. NPJ Aging Mech Dis 2, 16017 (2016).

43. C. R. Martens et al., Chronic nicotinamide riboside supplementation is well-tolerated and elevates NAD(+) in healthy middle-aged and older adults. Nature Communications 9, (2018).

44. J. P. van Dijken et al., An interlaboratory comparison of physiological and genetic properties of four Saccharomyces cerevisiae strains. Enzyme Microb Technol 26, 706–714 (2000).

45. T. Wang et al., Identification and characterization of essential genes in the human genome. Science 350, 1096–1101 (2015).

46. J. Schindelin et al., Fiji: an open-source platform for biological-image analysis. Nat Methods 9, 676–682 (2012).

47. A. M. Hosios et al., Amino Acids Rather than Glucose Account for the Majority of Cell Mass in Proliferating Mammalian Cells. Developmental cell 36, 540–549 (2016).

48. A. Panov, Z. Orynbayeva, Bioenergetic and Antiapoptotic Properties of Mitochondria from Cultured Human Prostate Cancer Cell Lines PC-3, DU145 and LNCaP. PlosOne 8, (2013).

49. C. Gregg, P. Kyryakov, V. I. Titorenko, Purification of mitochondria from yeast cells. J Vis Exp, (2009).

50. J. K. Salabei, A. A. Gibb, B. G. Hill, Comprehensive measurement of respiratory activity in permeabilized cells using extracellular flux analysis. Nature Protocols 9, 421–438 (2014).

51. F. Palmieri, C. L. Pierri, A. De Grassi, A. Nunes-Nesi, A. R. Fernie, Evolution, structure and function of mitochondrial carriers: a review with new insights. Plant J 66, 161–181 (2011).

52. R. C. Edgar, MUSCLE: multiple sequence alignment with high accuracy and high throughput. Nucleic Acids Res 32, 1792–1797 (2004).

53. J. Felsenstein, PHYLIP - Phylogeny Inference Package (Version 3.2). Cladistics 5, 164–166 (1989).

54. J. P. Ebejer, J. R. Hill, S. Kelm, J. Shi, C. M. Deane, Memoir: template-based structure prediction for membrane proteins. Nucleic Acids Res 41, W379–383 (2013).

55. E. F. Pettersen et al., UCSF Chimera--a visualization system for exploratory research and analysis. J Comput Chem 25, 1605–1612 (2004).

56. J. Barretina et al., The Cancer Cell Line Encyclopedia enables predictive modelling of anticancer drug sensitivity. Nature 483, 603–607 (2012).

